# RNA helicase DDX5 enables STAT1 mRNA translation and interferon signaling in hepatitis B virus replicating hepatocytes

**DOI:** 10.1101/2020.09.25.313684

**Authors:** Jiazeng Sun, Guanhui Wu, Wen-Hung Wang, Zhengtao Zhang, Philippe Merle, Lijian Hui, Danzhou Yang, Ourania Andrisani

## Abstract

**Objective:** RNA helicase DDX5 is downregulated during hepatitis B virus (HBV) replication, and poor prognosis HBV-related hepatocellular carcinoma (HCC). The objective of this study is to understand the significance of DDX5 downregulation during HBV infection and poor prognosis HCC, as it pertains to therapy of chronically HBV-infected patients with HCC. IFN-α therapy is effective only for a subgroup of HBV-infected patients, for reasons not understood. Herein, we investigated a novel mechanism of STAT1 translational control, involving DDX5-mediated resolution of a G-quadruplex structure, located at the 5’UTR of STAT1 mRNA, enabling STAT1 translation.

**Design and Results:** Molecular, pharmacologic, and biophysical assays were used together with cellular model of HBV replication, HCC cell lines, and HBV-related liver tumors. We observed, the protein level of DDX5 correlated with that of transcription factor STAT1. We show, DDX5 regulates STAT1 mRNA translation by resolving a G-quadruplex (rG4) RNA structure, proximal to the 5’ end of STAT1 5’UTR. We employed luciferase reporter assays comparing wild type (WT) *vs*. mutant (MT) rG4 sequence, rG4-stabilizing compounds, CRISPR/Cas9 editing of the rG4 sequence, and circular dichroism determination of rG4 structure. Ribonucleoprotein assays identified direct DDX5 binding to WT but not MT rG4. Importantly, liver cancer cell lines expressing low DDX5 exhibited reduced interferon response, while immunohistochemistry of HBV-related HCCs exhibiting absence of DDX5, also lacked STAT1.

**Conclusion:** DDX5 resolves a G-quadruplex structure in 5’UTR of STAT1 mRNA, enabling STAT1 translation. DDX5 is a key regulator of the dynamic range of interferon response during innate immunity and adjuvant IFN-α therapy.

## Introduction

Interferon Type I/III signaling acting through the JAK/STAT pathway exerts antiviral, antitumor, and immunomodulatory effects (1, 2). Effects of IFN-α and IFN-β are mediated by tyrosine phosphorylation of transcription factors STAT1 and STAT2, followed by formation of ISGF3 complex via interaction with IRF9, translocation to the nucleus, and transcriptional induction of Interferon Stimulated Genes (ISGs). In addition to tyrosine phosphorylation, STAT1 activity is further modulated by modifications including methylation (3), acetylation (4) and ubiquitination (5). Since STAT1 is involved in all types (I/III) of interferon signaling (6), these post-translational modifications influence all types (I/III) of interferon signaling, depending on cellular context. Herein, we provide evidence for a novel mechanism of STAT1 regulation, involving translational control of STAT1 mRNA, studied in the context of viral replication in hepatocytes harboring the hepatitis B virus (HBV).

Hepatitis B virus (HBV) infection is a significant global health problem with more than 250 million chronically infected patients. Chronic HBV infection is associated with progressive liver disease, cirrhosis, and development of hepatocellular carcinoma (HCC). Curative treatments for early stage HCC include liver resection, transplantation, or local ablation. In advanced stage HCC, multi-kinase inhibitors (7–9), offer only palliative benefits. In addition, because IFNs exert both antiviral and anti-proliferative effects (10, 11), adjuvant IFN-α therapy has been extensively applied for treatment of chronically HBV-infected patients with HCC. Unfortunately, many patients do not respond to IFN-α (12). Importantly, HCC patients with increased expression of the ISG IFIT3, a direct transcription target of STAT1 (13), predict positive response to IFN-α therapy (14), suggesting IFN-α non-responders lack STAT1 activation and/or expression.

Viruses have evolved effective strategies to hijack cellular pathways for their growth advantage, including mechanisms for immune evasion. In our studies, we have identified a cellular mechanism hijacked by HBV associated with both viral biosynthesis and poor prognosis HBV-related HCC (15–17). This mechanism involves the chromatin modifying PRC2 complex and RNA helicase DDX5. PRC2 mediates repressive histone modifications (H3K27me3)(18), while RNA helicase DDX5 is involved in transcription, epigenetic regulation, miRNA processing, mRNA splicing, decay, and translation (19, 20). Interestingly, HBV infection downregulates DDX5 via induction of two microRNA clusters (21), miR17~92 and miR106~25 (15–17). Moreover, HBV replicating cells with reduced DDX5 exhibit Wnt activation, resistance to chemotherapeutic agents (21), and reduced protein levels of STAT1, as we describe herein.

In this study, we provide evidence that DDX5 exerts additional functions in HBV infected cells, i.e., modulation of the innate immune response, via a novel mechanism of translational control of STAT1. Specifically, RNA helicase DDX5 resolves a secondary RNA structure called G-quadruplex (22), located in the 5’ untranslated region (5’UTR) of STAT1 mRNA, thereby enabling its translation. G-quadruplexes are four-stranded structures formed in guanine (G)-rich sequences (23), and when located in 5’UTRs of mRNAs, influence post-transcriptional regulation of gene expression (20). One of the first examples of translational repression by an RNA G-quadruplex (rG4) was demonstrated for human NRAS mRNA (24). Bioinformatics analysis estimated nearly 3,000 5’UTR rG4s in the human genome (24), and rG4-sequencing of polyA+ HeLa RNA generated a global map of thousands rG4 structures (25). Recently, it was discovered that DDX5 proficiently resolves both RNA and DNA G4 structures (26).

The significance of this mechanism of translational control of STAT1 is dependence on the protein level and activity of the rG4-resolving helicase. In terms of IFN (I/III) signaling and the innate response of HBV infected cells, our results presented herein demonstrate that the magnitude of the interferon response is influenced by this dynamic mechanism of STAT1 translational regulation, dependent on the protein level of DDX5.

## Materials and Methods

### Cell culture

Human hepatocellular carcinoma (HCC) cell lines HepG2, Huh7, Snu387, Snu423, HepaRG (27), HepAD38 (28), CLC15 and CLC46 (29) were grown as described, and regularly tested for mycoplasma using PCR Mycoplasma Detection Kit (Abm). HepAD38 cells support HBV replication by tetracycline removal (28). HepAD38 were STR tested by ATCC.

### Transfection assays

HepAD38, Huh7, and HepaRG cells (0.3×10^6^ cells per well of 6-well plate) were transfected with control (Ctrl) or DDX5 siRNAs (40 nM). Following transfection (48 h), cells were harvested for RNA or protein extraction and analyzed by qRT-PCR or immunoblotting, respectively. HepAD38, Huh7, and HepaRG cells (0.1×10^6^ cells per well of 12-well plate) were co-transfected with Renilla luciferase (25 ng) and WT STAT1-5’UTR-Luciferase (pFL-SV40-STAT1-5’UTR a gift from Dr. Ming-Chih Lai) (50 ng) or mutant MT-STAT1-5’UTR-Luciferase (50 ng) using Lipofectamine 3000 (Life Technologies). In HepAD38 cells, HBV replication was induced by tetracycline removal 4 days prior to transfection with Renilla and Firefly luciferase vectors. Ctrl or DDX5 siRNAs (40 nM each) were co-transfected with Renilla and Firefly luciferase vectors using RNAiMax (Life Technologies). Plasmids used are listed in Supporting Table S1. Firefly luciferase activity was measured 24 h after transfection using Dual Luciferase Assay system (Promega) according to manufacturer’s protocol, normalized to Renilla luciferase.

### RNA preparation and qRT-PCR assay

RNA preparation and qRT-PCR performed according to manufacturer’s instructions, employing commercially available kits (Supporting Table S2). Primer sequences listed in Supporting Table S3.

### Immunoblot and Immunohistochemistry assays

performed as described (16). Densitometric analysis of immunoblots was by ImageJ. Antibodies used are listed in Supporting Table S4.

### Ribonucleoprotein Immunoprecipitation (RIP) Assay

RIP assays employed the Magna RIP™ RNA-Binding Protein Immunoprecipitation Kit (Millipore Sigma) following manufacturer’s instructions. Primer sequences and antibodies used are listed in Supporting Table S3 and S4, respectively.

### RNA pull down assay

RNA folding and pull down assays were performed as described with modifications (30). Briefly, synthetic 5’-biotinylated rG4 (Bio-rG4) and rG4mut (Bio-rG4mut) RNA oligonucleotides (Millipore Sigma) were diluted to 5 mM in folding buffer, heated to 95 °C, and cooled to 25°C. G-quadruplex formation determined by circular dichroism (CD), as described below. Whole cell extracts from HepAD38, Huh7, and HepaRG cells were incubated with folded biotinylated RNAs (Table S3), followed by pull down with streptavidin beads (Promega). Bound proteins analyzed by immunoblotting.

### Circular Dichroism spectroscopy

of RNA oligonucleotides performed as described (31), employing Jasco J-1100 spectropolarimeter equipped with thermoelectrically controlled cell holder. CD measurements were performed using quartz cell with optical path length of 1 mm. Blank sample contained only buffer (25mM Tris-HCL, pH 7.4). Each CD spectroscopy measurement was the average of two scans, collected between 340 and 200 nm at 25°C, scanning speed 50 nm/min. CD melting experiments were performed at 264 nm with heating rate of 2°C/min between 25°C and 95°C. RNA sample concentration was 10 μM.

### CRISPR/Cas9 gene editing

Huh7 and HepaRG cells were used to introduce indels targeting essential nucleotides of the G-quadruplex structure in 5’ UTR region of DDX5 gene (32). Ribonucleoprotein (RNP) of Cas9-2NLS (10 pmol) and guide RNA (50 pmol, Synthego) were loaded onto a 10 μl Neon Tip, and electroporated into 1×10^5^ Huh7 and HepaRG cells, using Neon Transfection System at 1200 V, for 20 msec and 4 pulses (ThermoFisher Scientific), according to manufacturer’s instructions. Genomic DNA was isolated and used for rapid PAGE genotyping (33) to validate incorporation of indels, 48 h after electroporation. Validated pools of cells were subjected to clonal selection. Isolated single colonies were confirmed by rapid PAGE genotyping and allelic sequencing.

### Statistical Analysis

Statistical analysis was performed using unpaired *t* test in GraphPad Prism version 6.0 (GraphPad Software, San Diego, CA). Differences were considered significant when *p* < 0.05.

## Results

We examined various liver cancer cell lines, HepG2, Huh7, Snu423 and HepaRG for IFN-α response, employing immunoblots for activated phospho-STAT1 (T701) and expression of ISG IRF9. All cell lines responded to IFN-α, including the HBV replicating HepAD38 cell line that contains a stably integrated copy of the HBV genome under control of the Tet-off promoter (28) (Figure 1A and Supplementary Figure S1A-B). In the absence of IFN-α treatment, HBV replication in HepAD38 cells exerted a small but reproducible reduction on STAT1 protein level (Figure 1A). Following IFN-α treatment for 24 h, levels of STAT1, p-STAT1, and downstream ISGs IRF9 and IFTM3 were similarly reduced (Figure 1A and Supplementary Figure S1C). Likewise, HBV replication exerted a reproducible decrease in protein level of DDX5, irrespective of IFN-α treatment (Figure 1A), as reported previously (16).

**Figure 1.**
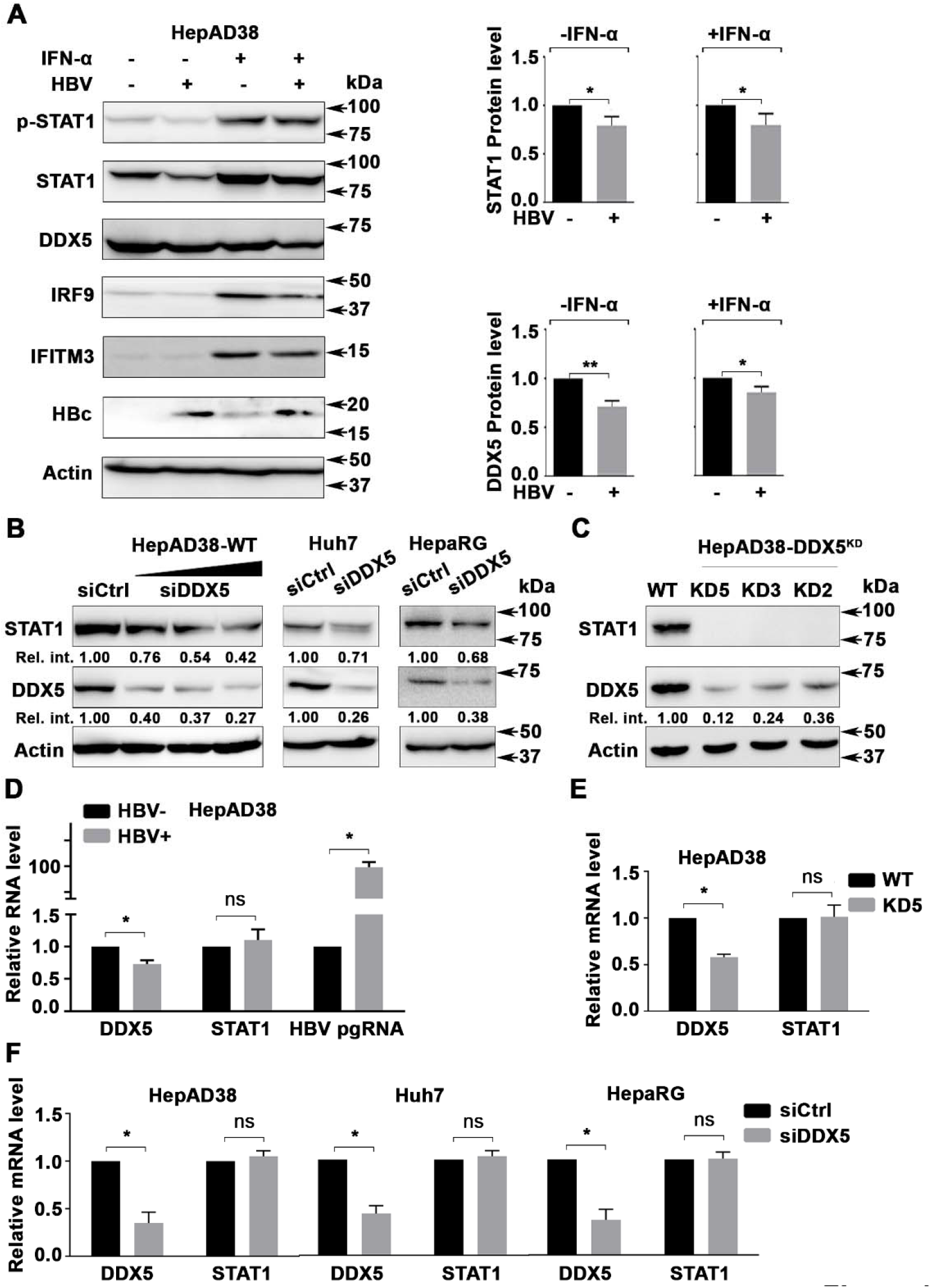
DDX5 knockdown regulates STAT1 mRNA translation. **(A)** Immunoblots of IFN-α induced proteins using lysates from HepAD38 cells with (+) or without (-) HBV replication for 5 days as a function of IFN-α (500 ng/ml) treatment for the last 24 h. (Right panel) quantification of DDX5 and STAT1 protein level, by ImageJ software, from three independent biological replicates. *: p<0.05, **: p<0.01; Error bars indicate Mean ±SEM. Immunoblot of indicated proteins in: **(B)** HepAD38, Huh7 and HepaRG cells transfected with DDX5 siRNA (siDDX5) or negative control siRNA (siCtrl) for 48 h, and **(C)** in WT and DDX5 knockdown HepAD38 cell lines KD2, KD3 and KD5. **(D)-(F)** qRT-PCR of HBV pgRNA and STAT1 mRNA using RNA from **(D)** HepAD38 cells with (+) or without (-) HBV replication for 5 days; **(E)** WT and KD5 HepAD38 cells, and **(F)** HepAD38, Huh7, and HepaRG cells transfected with siDDX5 or siCtrl for 48h. Statistical analysis of DDX5 and STAT1 mRNA levels from three biological replicates. *: p<0.05, ns: not significant; Error bars indicate Mean ±SEM.

To determine whether DDX5 downregulation was associated with downregulation of STAT1 observed during HBV replication (Figure 1A), we transfected increasing amounts of siRNA for DDX5 in HepAD38 cells. Surprisingly, reduction in DDX5 protein resulted in progressive reduction in STAT1 protein level (Figure 1B and Supplementary Figure S2). DDX5 knockdown in Huh7 and HepaRG cell lines also resulted in reduced STAT1 protein (Figure 1B and Supplementary Figure S2), whose t1/2 was quantified to be 16 h (Supplementary Figure S3A-B). Next, we examined STAT1 protein levels in three clonal DDX5-knockdown cell lines, KD2, KD3 and KD5, constructed in HepAD38 cells. These cell lines lacked STAT1 protein (Figure 1C) and I IFN-α response (Supplementary Figure S3C). Interestingly, STAT1 mRNA levels were unaffected in HBV replicating HepAD38 cells (Figure 1D), in DDX5 knockdown HepAD38 cells (Figure 1E), and upon transient siRNA-mediated DDX5 knockdown in HepAD38, Huh7, and HepaRG cells (Figure 1F), thereby excluding DDX5 effects on STAT1 transcription.

The STAT1 gene contains 24 introns. Accordingly, we examined whether intron detention or aberrant splicing are regulated by DDX5. qRT-PCR and RNA-seq intron data analyses of STAT1 in WT HepAD38 cells *vs*. DDX5-knockdown cells excluded STAT1 intron detention (Supplementary Figure S4A). Likewise, comparison of STAT1 mRNA sequence from WT *vs*. DDX5-knockdown cells, excluded aberrant splicing involving the mRNA splice site in proximity to AUG (Supplementary Figure S4B). These results suggested DDX5 regulates STAT1 mRNA translation.

### rG4 structures in 5’UTR of human STAT1 mRNA

Having excluded DDX5 effects on STAT1 mRNA transcription and processing, we reasoned, the information for post-transcriptional regulation must be located in the sequence or structure of STAT1 5’ UTR. Surprisingly, the rG4-seq transcriptomic studies by Kwok et al (25) identified the 5’UTR of human STAT1 mRNA as a high probability mRNA harboring rG4 structures (Supplementary Figure S4C). To test this hypothesis, and using NRAS as our positive control (24), we examined the effect of several G4 stabilizing compounds (Figure 2A), including PhenDC3 and RR82 (22), on STAT1 protein level. Both compounds reduced STAT1 and NRAS protein level, without affecting STAT1 mRNA (Figure 2B). Similarly, we tested the effect of the G4-interactive TMPyP4 and its corresponding non-G4-interactive TMPyP2 compound (34) on STAT1 protein and mRNA levels, using HepAD38 and Huh7 cells (Figure 2C). The G4-interactive TMPyP4, similar to RR82, suppressed STAT1 protein levels, whereas the non-G4 interactive TMPyP2 exerted no effect. Importantly, these compounds did not affect STAT1 mRNA levels (Figure 2C). We interpret these results to mean the 5’UTR of STAT1 contains a putative rG4 structure, regulating STAT1 expression post-transcriptionally.

**Figure 2.**
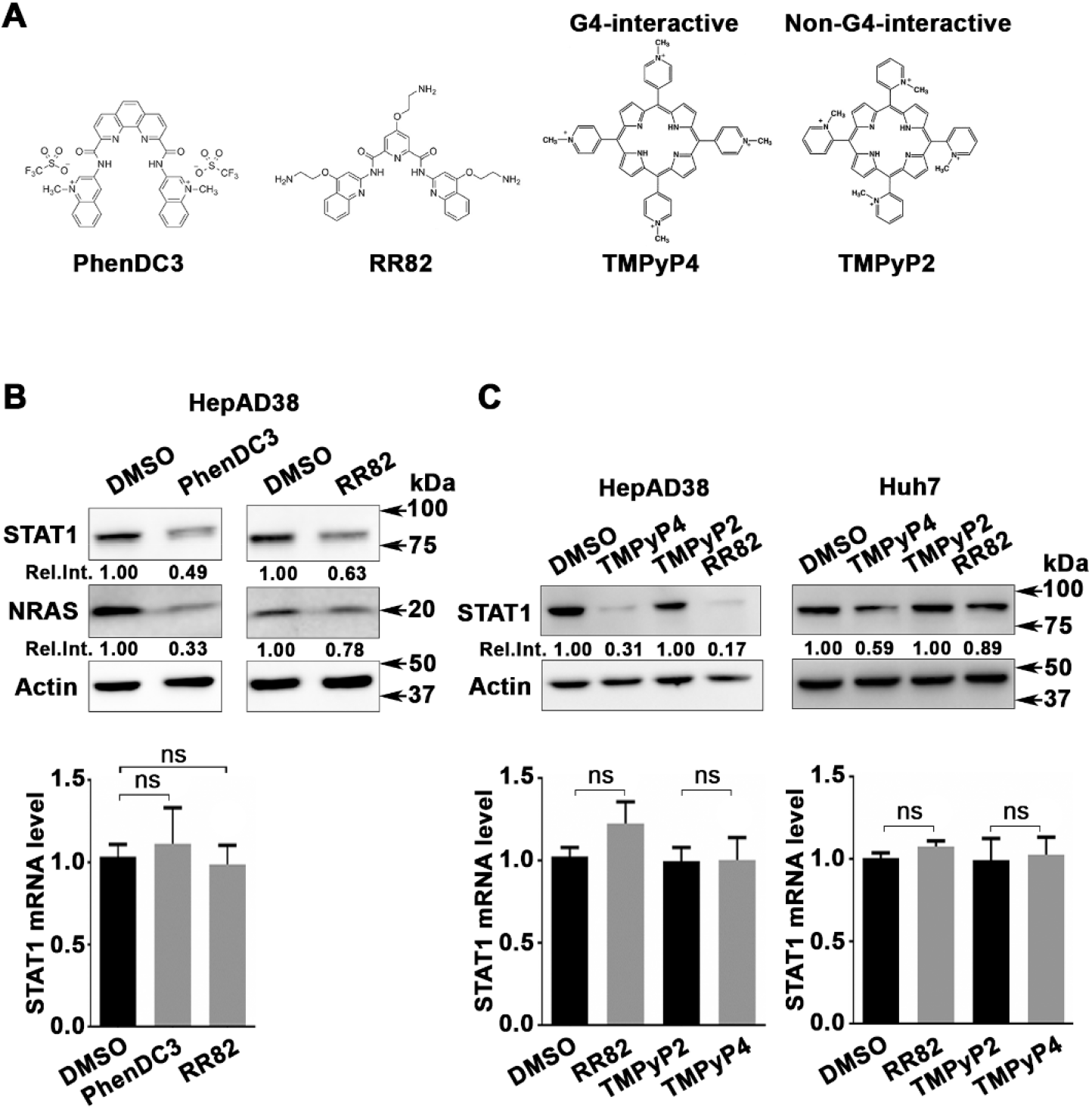
G-quadruplex stabilizing drugs reduce STAT1 protein levels. **(A)** Chemical structure of G4 stabilizing compounds PhenDC3, RR82, TMPyP4 and TMPyP2. **(B)** Immunoblots of STAT1 and NRAS, using lysates from HepAD38 cells treated for 48 h with PhenDC3 (5 μM) or RR82 (5 μM). (Lower panel) qRT-PCR of STAT1 mRNA using RNA from HepAD38 cells treated with DMSO, PhenDC3 (5 μM) or RR82 (5 μM) for 48 h. **(C)** Immunoblots of STAT1 from lysates of HepAD38 and Huh7 cells treated with TMPyP4 (5 μM) or TMPyP2 (5 μM) for 48h. (Lower panel), qRT-PCR of STAT1 mRNA from HepAD38 and Huh7 cells, treated as indicated. Statistical analysis of STAT1 mRNA is from three biological replicates. ns: not significant; Error bars indicate Mean ±SEM.

Employing an expression vector driven by the SV40 promoter, we cloned nucleotides (nt) + 1 to +400 of the 5’UTR of STAT1 upstream of Firefly (F.) luciferase gene. This 5’UTR region contains three putative rG4s, labeled as rG4-1, rG4-2 and rG4-3 (Supplementary Figure S4C). Importantly, rG4-1 is located 30 nt downstream from the start of the 5’UTR. Earlier studies demonstrated rG4s situated proximal or within the first 50 nt to the 5’ end of the 5’UTR are functional and effective in repressing translation (35). Based on this reasoning, we focused our analysis on rG4-1 and constructed a mutant (MT) rG4-1 (Figure 3A), with G to A substitutions within the putative rG4-1 sequence. Transfection in HepAD38 cells of expression vectors containing WT and MT rG4-1 demonstrated a statistically significant increased F. luciferase activity from MT rG4-1 in comparison to WT rG4-1 vector, while no changes were observed at the mRNA level, both in HepAD38 and Huh7 cells (Figure 3B). Mutational analyses of rG4-2 and rG4-3 sequences excluded a similar role on STAT1 expression (Supplementary Figure S5A and B). Next, we examined the effect of HBV replication, using HepAD38 cells, on F. luciferase activity expressed from WT and MT rG4-1 vectors. HBV replication reduced F. luciferase activity only from the WT rG4-1 containing vector, without an effect on F. luciferase mRNA expression (Figure 3C). Similar results were observed by siRNA knockdown of DDX5 (siDDX5) (Figure 3C). Employing these expression vectors, we also tested the effect of the G4-stabilizing compounds RR82 and TMPyP4. Both drugs reduced expression only from the WT rG4-1 vector, without an effect on mRNA levels of F. luciferase, tested in HepAD38 (Figure 3D) and Huh7 (Figure 3C) cells. Taken together, these results identify the rG4-1 sequence as a functional element in 5’UTR of STAT1 mRNA, regulating its translation.

**Figure 3.**
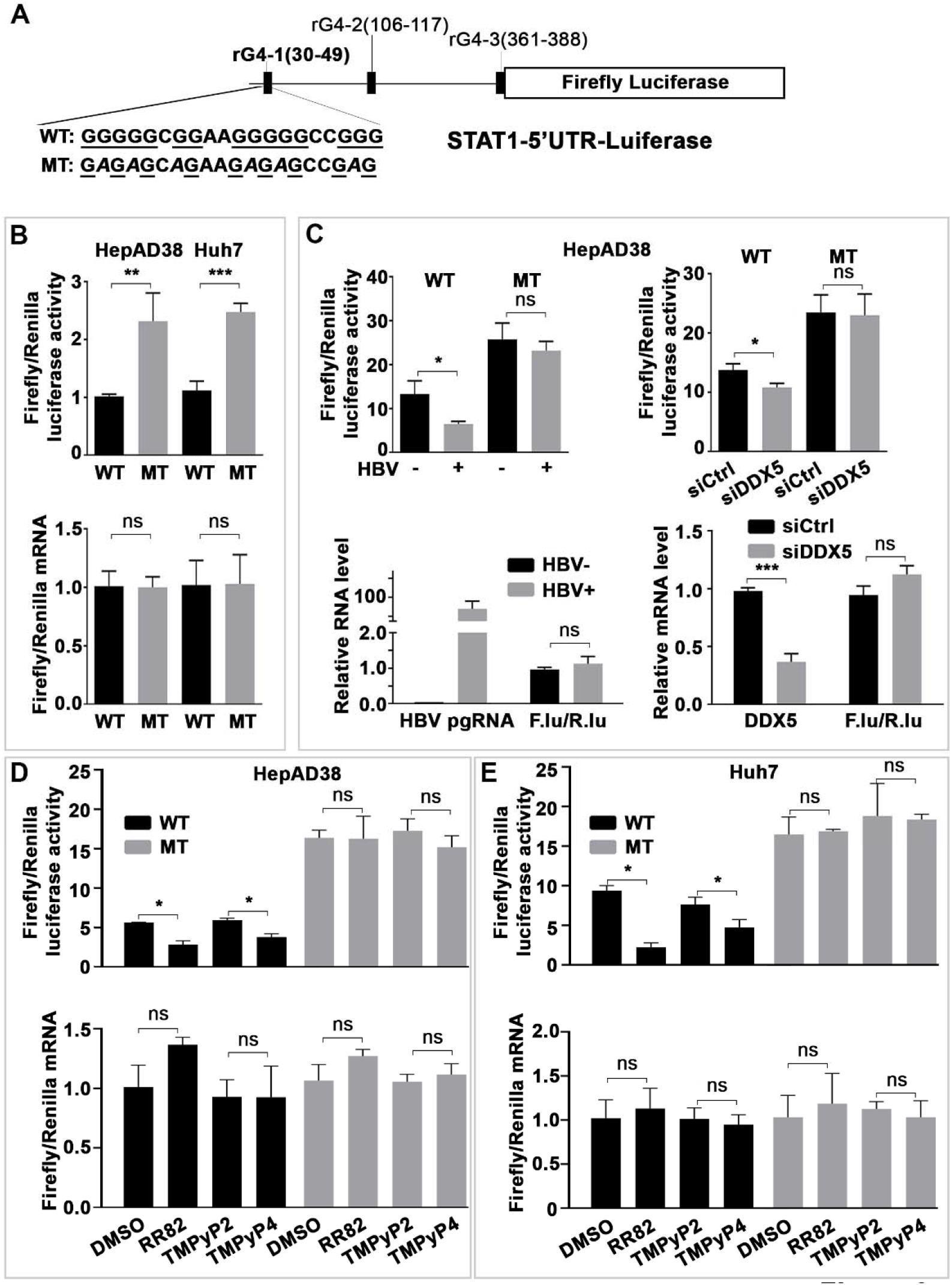
G-quadruplex (rG4) regulates STAT1 expression post-transcriptionally. **(A)** Human STAT1 5’UTR upstream of Firefly (F.) Luciferase reporter, driven from SV40 promoter. Putative rG4 sequences in 5’ UTR indicated as rG4-1, rG4-2 and rG4-3. WT rG4-1 nucleotide sequence is shown. Italics indicate site-directed changes in mutant MT-rG4-1. **(B)** - **(E)** Ratio of Firefly/Renilla luciferase activity at 24 h after co-transfection of WT or MT STAT1-5’UTR-F. Luciferase and Renilla-Luciferase expression plasmids. Lower panels, ratio of Firefly/Renilla luciferase mRNAs quantified by qRT-PCR. **(B)** HepAD38 and Huh7 cells. **(C)** HepAD38 cells with (+) or without (-) HBV replication for 5 days. **(D)** HepAD38 cells, and **(E)** Huh7 cells treated with indicated G-quadruplex stabilizing drugs (5 μM) for 24 h. Statistical analysis from three independent biological replicates. *: p<0.05, **: p<0.01, ***: p<0.001, ns: not significant; Error bars indicate Mean ±SEM.

### Genomic editing of rG4-1 increases STAT1 protein levels

Using clustered regularly interspaced short palindromic repeats (CRISPR)/Cas9 technology (36) we edited the genomic rG4-1 sequence of STAT1 in Huh7 and HepaRG cell lines. Several clones were isolated and sequenced, and protein and mRNA levels of STAT1 were determined (Figure 4 and Supplementary Figure S6). DNA sequencing of HepaRG clones C5 and C9 (Supplementary Figure S6) demonstrated C5 cells contain rG4-1 deletions in both alleles, while C9 cells have rG4 changes only in one allele (Figure 4B). All clones analyzed from Huh7 and HepaRG cells exhibited statistically significant and reproducible increases in protein level of STAT1 in comparison to WT (unedited) cells, while STAT1 mRNA levels remained unchanged (Figure 4). These results are also supported by luciferase reporter assays containing the edited rG4-1 sequences (Supplementary Figure S7).

**Figure 4.**
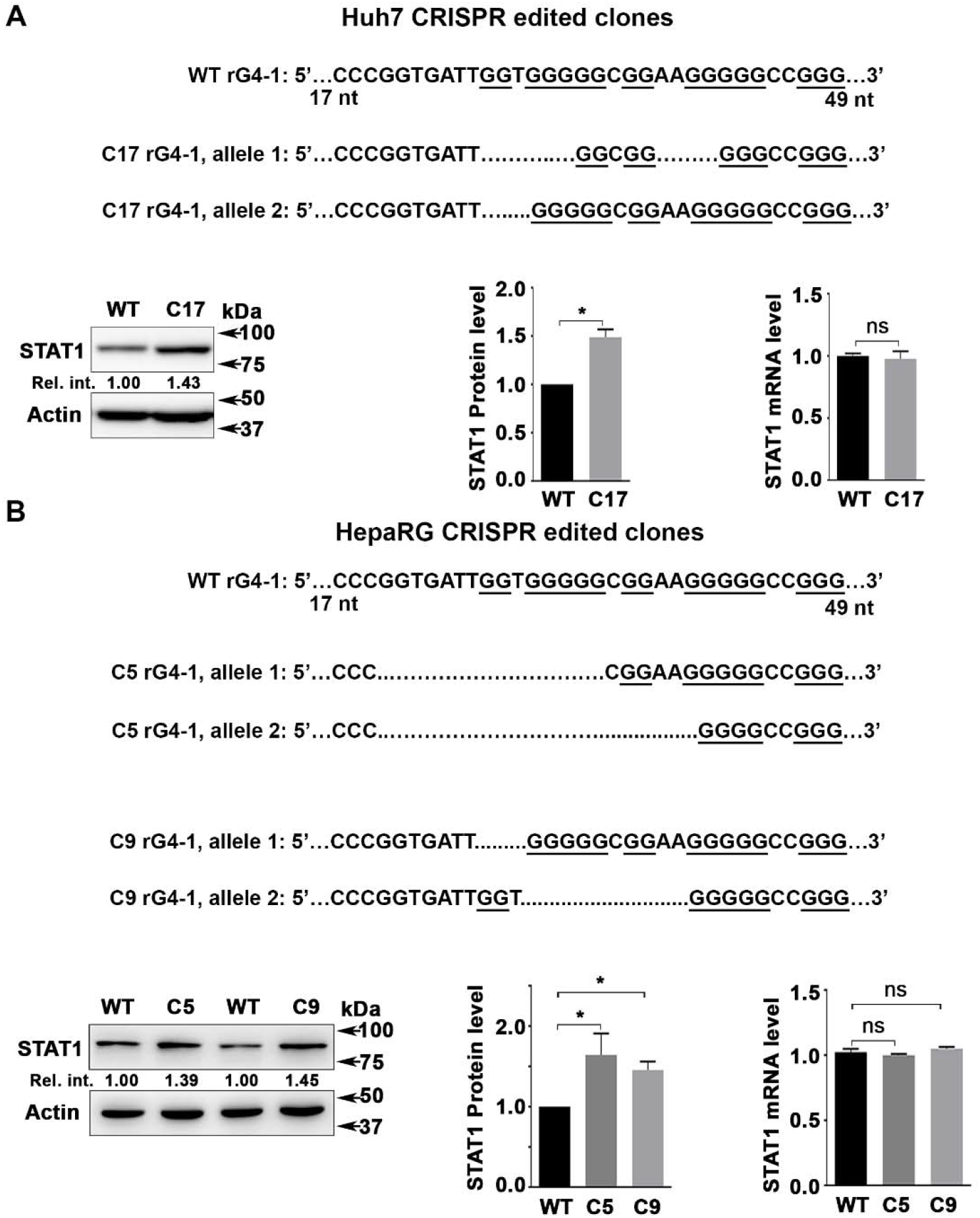
Genomic editing of rG4-1 increases STAT1 protein levels. **(A)** Sequence of CRIPSR/Cas9 edited rG4-1 sequence in **(A)** Huh7 and **(B)** HepaRG cells. Immunoblot of STAT1 in indicated cell lines. Right panels, quantification of STAT1 protein by ImageJ software, and qRT-PCR of STAT1 mRNA, from three independent biological replicates. *: p<0.05, ns: not significant; Error bars indicate Mean ±SEM.

Interestingly, DDX5 knockdown by transfection of DDX5 siRNA had no effect on STAT1 protein levels of clone C5, containing edited rG4 sequence on both STAT1 alleles (Figure 5A). By contrast, siDDX5 reduced STAT1 protein levels in WT and C9 cells (Figure 5A). Likewise, STAT1 protein levels of clone C5 were resistant to G4-stabilizing drugs RR82 and TMPyP4, while these compounds reduced STAT1 levels in both WT and C9 cells (Figure 5B). Next, we examined the interferon response of WT, C5, and C9 cells as a function of cotreatment with RR82 (Figure 5C). In contrast to WT and C9 cells, the IFN-α response of C5 cells was not inhibited by RR82, in terms of STAT1 protein level and activation, as well as induction of IRF9 and IFITM3 (Figure 5C and Supplementary Figure S8). These results support that both alleles of clone C5 contain nonfunctional rG4-1 structures in 5’UTR of STAT1 mRNA.

**Figure 5.**
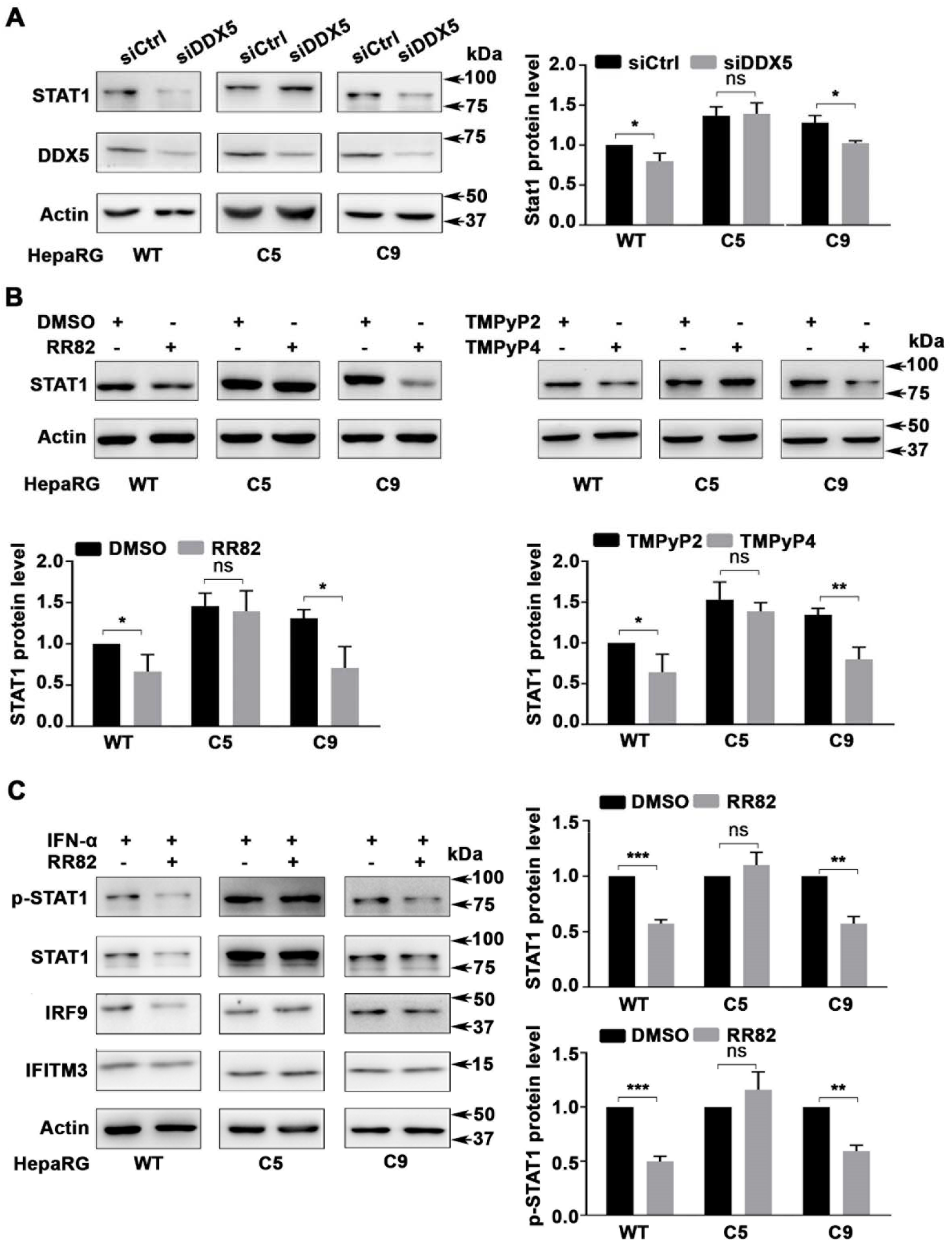
Effect of G-quadruplex stabilizing drugs on rG4-1 edited HepaRG cells. Immunoblots of indicated proteins using lysates from indicated HepaRG cells (WT, C5, and C9). **(A)** After transfection of siCtrl or siDDX5 RNA for 48 h; **(B)** following treatment with RR82 (5 μM), TMPyP4 (5 μM) or TMPyP2 (5 μM) for 48 h, and **(C)** treatment with RR82 (5 μM) for 48 h in combination with IFN-α (500 ng/ml) for the last 24 h. Quantification **(A-C)** from three independent biological replicates. *: p<0.05, **: p<0.01, ***: p<0.001, ns: not significant. Error bars indicate Mean ±SEM.

### The rG4-1 sequence of STAT1 forms G-quadruplex *in vitro*

To directly determine whether the rG4-1 sequence in 5’UTR of STAT1 indeed forms a G-quadruplex structure, we employed CD spectroscopy, a method that enables study of nucleic acid secondary structure (31, 37). RNA oligonucleotides were synthesized (Figure 6A) that corresponded to the indicated sequences of rG4-1 found in WT STAT1 mRNA, and in each allele of clones C5 and C9. Since G4 oligonucleotides can form higher-order structures, we included U-streches at the 5’ and 3’ of the indicated RNA oligonucleotides, and confirmed these RNA molecules were in monomeric form, employing native PAGE (Supplementary Figure S9).

**Figure 6.**
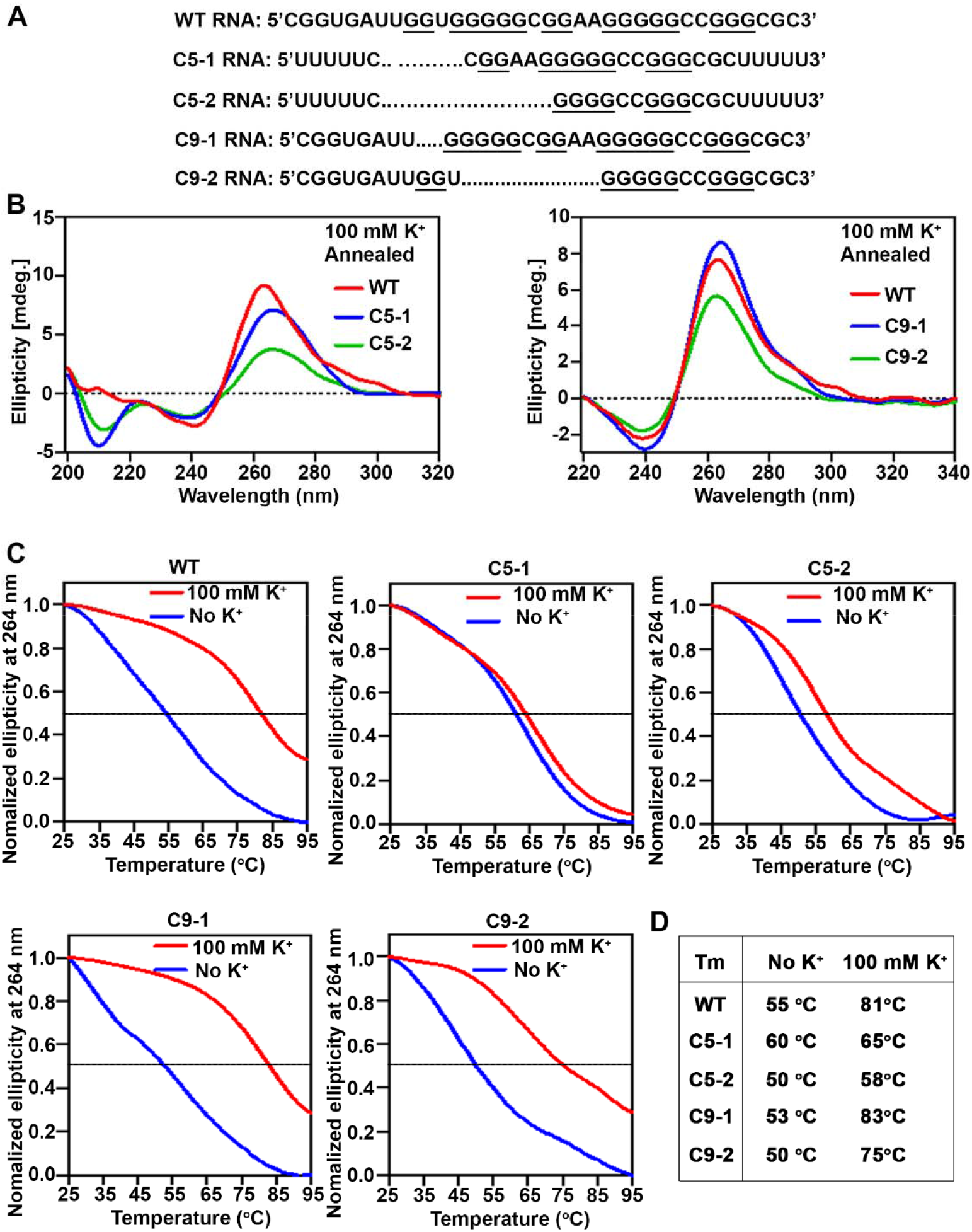
The rG4-1 sequence in 5’UTR of STAT1 mRNA forms G-quadruplex. **(A)** Synthetic RNA oligonucleotides of WT STAT1 rG4-1 and corresponding mutants in HepaRG clones C5 and C9. **(B)** CD spectroscopy measurement, and **(C)** melting curves of RNA oligonucleotides annealed by heating to 95°C and slowly cooled down to room temperature. **(D)** Melting temperature (Tm), without and with (100 mM) KCl, calculated from **A-C.**

The CD spectrum and thermal stability of these RNA oligonucleotides were studied as a function of K+ addition (100 mM), a cation that stabilizes G4 structures (38). The melting temperature (Tm) of each RNA oligonucleotide is shown (Figure 6D). Addition of 100 mM KCl increased the Tm of the WT, C9-1, and C9-2 RNA oligonucleotides, supporting formation of stable rG4 structures. By contrast, oligonucleotide C5-1 and C5-2 exhibited no or very small Tm increase in comparison to WT. These results indicate, the WT, C9-1 and C9-2 sequences form stable rG4 structures, whereas C5-1 and C5-2 lack formation of this structure. These biophysical results are congruent with the biological data presented herein, and demonstrate the role of rG4-1 structure in STAT1 translational control.

### DDX5 selectively binds STAT1 mRNA with WT rG4-1

To determine whether DDX5 interacts directly with STAT1 mRNA, we performed ribonucleoprotein immunoprecipitation (RIP) assays employing DDX5 antibody and IgG as negative control. In HepAD38 and Huh7 cells endogenous, immunoprecipitated DDX5 was found in association with STAT1 mRNA, while IgG did not exhibit such association (Figure 7A and B). Next, we performed RIP assays in HepaRG cells, WT and clones C5 and C9. Interestingly, endogenous DDX5 associated with STAT1 mRNA in WT and clone C9 cells, while STAT1 mRNA expressed in clone C5, containing mutated rG4-1 sequence on both alleles, lacked binding to DDX5 (Figure 7C).

**Figure 7.**
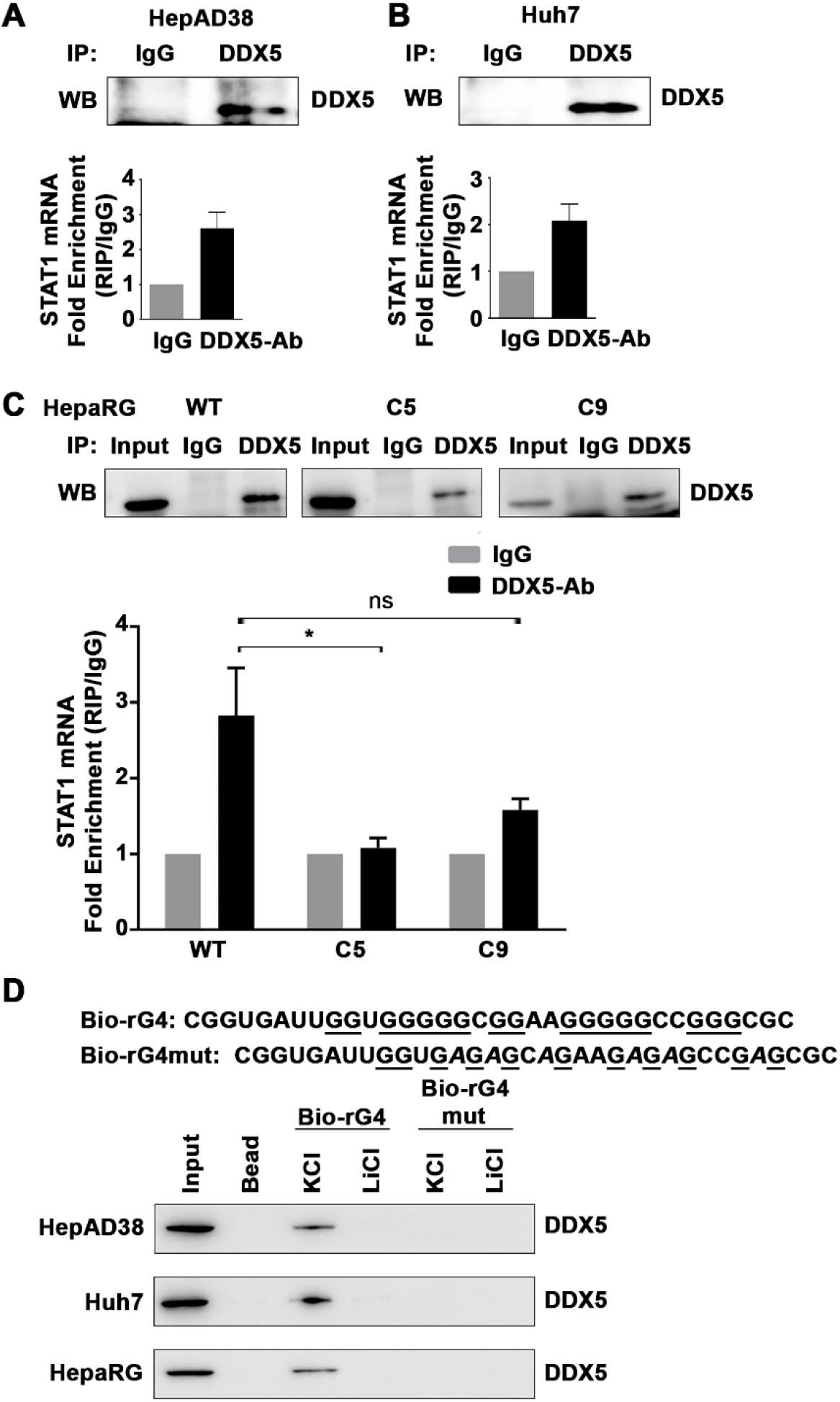
DDX5 binds STAT1 mRNA. Ribonucleoprotein immunoprecipitation (RIP) assays with DDX5 antibody performed in **(A)** HepAD38, **(B)** Huh7, and **(C)** HepaRG cells, WT, C5 and C9. DDX5 enriched RNAs quantified by qRT-PCR using STAT1 primers. Results are from three biological replicates. *: p<0.05, ns=not significant. **(D)** RNA pulldown assays using synthetic, biotinylated RNA oligonucleotides, WT and mutated rG4-1, in 100 mM KCl or 100 mM LiCl, bound to lysates from indicated cell lines, followed by immunoblots with DDX5 antibody. A representative assay is shown from three independent experiments.

To further confirm that DDX5 binds the rG4-1 sequence, biotinylated oligonucleotides containing WT or mutant rG4-1 (Figure 7D) were incubated in 100 mM K+ that stabilizes the G4 structure or 100 mM Li+ that does not (38). Mutant rG4-1 does not form stable G4 structure even in the presence of 100 mM K+, determined by CD spectroscopy (Supplementary Figure S10).

Next, pre-folded RNA oligonucleotides incubated with lysates from HepAD38, Huh7, and HepaRG cells were bound to streptavidin beads, and the retained proteins analyzed by DDX5 immunoblots. Indeed, these pull-down assays show DDX5 selectively binds the WT rG4-1 whose structure is stabilized by 100 mM K+ (Figure 7D).

### STAT1 expression in liver cancer cell lines and HBV-related liver tumors

To further establish the biological connection between DDX5 and STAT1 protein levels relative to interferon response, we quantified IFN-α response in liver cancer cell lines exhibiting different levels of DDX5 protein. We compared cell lines Snu387 *vs*. Snu423, and CLC15 *vs*. CLC46 (29), by performing immunoblots of lysates treated with increasing amount of IFN-α for 12h (Figure 8A and Supplementary Figure S11). Quantification of the immunoblots shows the p-STAT1, STAT1, DDX5 and IRF9 signal normalized to the baseline p-STAT1 signal obtained in the absence of IFN-α (Figure 8B). The results demonstrate, low level of DDX5 correlates with reduced STAT1 protein, STAT1 activation, and response to IFN-α. Likewise, we quantified the STAT1 mRNA from each cell line as a function of IFN-α, and normalized STAT1 protein levels quantified from the immunoblots (Figure 8A) relative to STAT1 mRNA. In comparison to Snu387 cells, Snu423 cells that express higher DDX5 protein also exhibit higher STAT1 protein (Figure 8C). Furthermore, we analyzed by immunohistochemistry the expression levels of DDX5 and STAT1 proteins in a small set of HBV-related HCCs. Employing normal human tissues as positive control, we observed absence of immunostaining for both DDX5 and STAT1 in Edmonson’s grade 3 HBV-related HCC tumors (Figure 8D), supporting our mechanistic *in vitro* observations.

**Figure 8.**
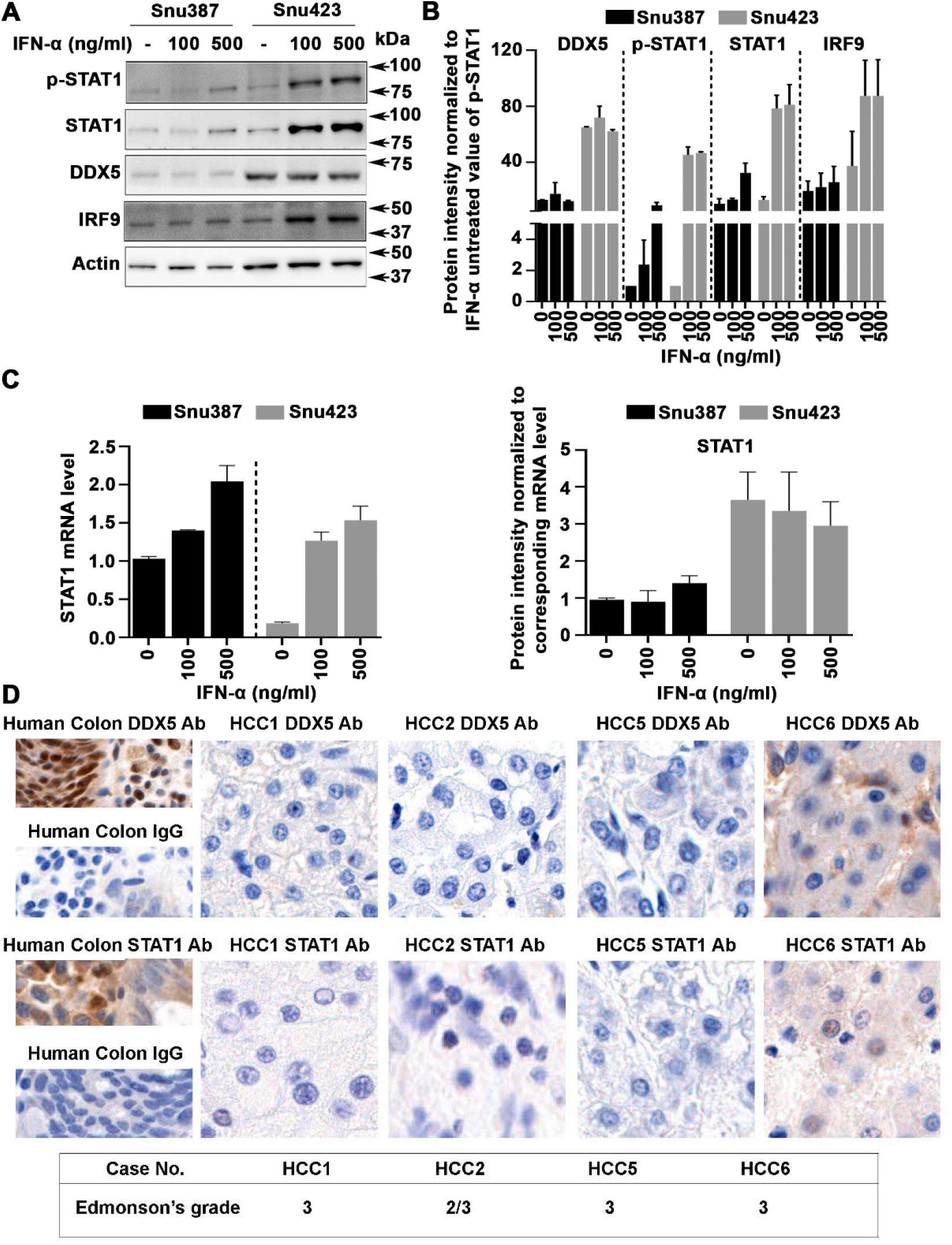
DDX5 expression level vs. magnitude of interferon response. **(A)** Immunoblots using lysates from indicated cell lines treated with IFN-α (100 and 500 ng/ml) for 12 h. **(B)** Quantification shows ratio of p-STAT1, STAT1, IRF9 and DDX5 relative to level of p-STAT1 in IFN-α untreated cells. Results are average from three independent experiments. **(C)**(Left panel) Quantification by RT qPCR of STAT1 mRNA in Snu387 and 423 cell lines treated with IFN-α as indicated for 12 h. (Right panel) STAT1 protein from **(A)** normalized to STAT1 mRNA in Snu387 *vs*. Snu423 cell lines, under the indicated treatment. **(D)** Immunohistrochemistry of HBV-related HCCs and normal human colon tissue, used as positive control, with DDX5 and STAT1 antibodies *vs*. IgG, performed as described (16).

## Discussion

Herein, we report the translational control of STAT1, a transcription factor essential for all types (I/III) of interferon signaling, via a G-quadruplex (rG4) structure located at the 5’UTR of STAT1 mRNA. RNA helicase DDX5 resolves this rG4 structure, thereby enabling STAT1 translation. Significantly, this post-transcriptional regulation of STAT1 expression is a cell intrinsic mechanism that influences the dynamic range of interferon response, dependent on protein level and activity of DDX5. The RNA helicase DDX5 regulates every aspect of RNA metabolism (39), and also serves as a barrier to pluripotency (40). Interestingly, it has been noted that pluripotent stem cells are refractory to interferon signaling (41, 42), although the mechanism is not yet understood. It is currently unknown whether absence of DDX5 expression or activity contributes to lack of innate immune response in pluripotent stem cells (41, 42).

G-quadruplexes (G4) are non-canonical, secondary DNA or RNA structures formed by guanine rich sequences (22, 23, 43). DNA G4 structures are found in gene promoters, particularly of oncogenes, regulating their transcription (44). Interestingly, recent studies demonstrated that DDX5 resolves both RNA and DNA G4 structures, including a G4 structure found in the MYC promoter, enhancing its transcriptional activity (26). RNA G-quadruplexes (rG4) are primarily located in 5’UTR or 3’UTR of mRNAs, playing important roles in RNA biology, from splicing, stability, mRNA targeting, and translation (22, 43). rG4 structures have been identified by various methods in the 5’ UTR of NRAS (24), Zic-1(45), Nkx2-5 (46) mRNAs, among other genes (22, 43), regulating their translation. Herein, we demonstrate by pharmacologic, molecular, and biophysical approaches the functional significance of the rG4 structure in the 5’UTR of STAT1 mRNA. Specifically, G4-structure stabilizing compounds RR82, PhenDC3 and TMPyP4 reduced the protein level of both STAT1 and NRAS, used as positive control, without affecting STAT1 mRNA level (Figure 2). After excluding other post-transcriptional modes of STAT1 regulation, we focused on the rG4-1 sequence located in proximity to the 5’ end of STAT1 5’UTR, since it had been identified as a high probability rG4 sequence by the transcriptomic study of Kwok et al (25). Indeed, using an SV40-driven Luciferase reporter, containing the STAT1 5’UTR (nt +1 to +400) upstream from F. Luciferase, we demonstrated that G4-structure stabilizing compounds suppressed the protein synthesis but not mRNA level of the WT rG4-1 containing luciferase vector (Figure 3). We also edited the genome of Huh7 and HepaRG cells by the CRISPR/Cas9 approach and generated alterations (deletions) of the rG4-1 sequence of STAT1 gene (Figure 4). When both alleles harbored the edited rG4-1 sequence, as in HepaRG clone C5, STAT1 protein levels were increased. Significantly, expression of STAT1 in clone C5 was resistant both to DDX5 knockdown (Figure 5A), and the inhibitory effect of G4-stabilizing compounds (Figure 5B). Circular dichroism and thermal stability measurements directly demonstrated formation of G-quadruplex only by WT and C9 rG4-1 sequences (Figure 6), supporting the biological data. RIP assays using antibody to endogenous DDX5 protein, showed lack of DDX5 binding to STAT1 mRNA from clone C5, containing edited rG4-1 sequence in both alleles. By contrast, DDX5 bound to STAT1 mRNA encoding the WT and C9 rG4-1 sequence (Figure 7). These results were further verified by RNA pulldown assays using synthetic RNA oligonucleotides under K+ conditions enabling formation of the G-quadruplex structure, thereby confirming endogenous DDX5 recognizes the G-quadruplex found at the 5’ UTR of STAT1 mRNA (Figure 7). Taken these results together, we conclude, the rG4-1 sequence in 5’UTR of human STAT1 mRNA assumes a G-quadruplex secondary structure. This rG4 structure, stabilized by G4-stabilizing compounds RR82, PhenDC3 and TMPyP4, suppresses STAT1 mRNA translation. RNA helicase DDX5 resolves the rG4 structure, enabling STAT1 mRNA translation.

Regarding the biological significance of this mechanism, we compared STAT1 levels and the interferon response using two sets of liver cancer cell lines expressing low *vs*. high DDX5, namely, Snu387 *vs*. Snu423, and CCL15 and CCL46 derived from HBV-related HCCs (29). We observed, the magnitude of the IFN-α response is a cell intrinsic property, dependent on the DDX5 expression level (Figure 8A-C and Supplementary Figure S11). Furthermore, although DDX5 is an ATPase (39), and arginine methylation by PRMT5 modulates the function of DDX5 in resolving DNA:RNA hybrids (47), how the activity of DDX5 is regulated during different cellular contexts, e.g., in response to IFNs, is presently not understood. Certainly, the regulation of DDX5 will add another level of complexity to this novel mechanism of interferon response.

Lastly, we show that in advanced HBV-related HCCs, there is absence both of DDX5 and STAT1 immunostaining (Figure 8D). Absence of STAT1 expression could provide an explanation for the lack of IFN-α response observed in many chronically HBV infected patients with HCC (14). Interestingly, a positive indicator for IFN-α responsiveness in HBV-infected patients with HCC (14) is expression of IFIT3, a STAT1 regulated gene (13). In addition, it is presently unknown whether this mechanism of DDX5-dependent STAT1 translational regulation is also linked to the immunotherapy response of patients with HCCs. Our earlier studies identified downregulation of DDX5 to be associated with poor prognosis HBV-related HCC (16, 48), and hepatocyte reprogramming to a less-differentiated state exhibiting features of hCSCs (21), in agreement with the role of DDX5 as a barrier to pluripotency (40). Regarding immunotherapy which is based on PD-1 blockade, the PD-L1,2 ligands are induced in tumors by interferon gamma, leading to immune evasion (49). According to the mechanism described herein, we propose that absence or downregulation of DDX5 in HCCs will result in absence or reduced protein of STAT1, and in turn reduced PD-L1 expression, rendering these tumors refractory to PD-1 targeting immunotherapy (49).

