## Supplementary Information for "RNA helicase DDX5 enables STAT1 mRNA translation and interferon signaling in hepatitis B virus replicating hepatocytes"

^1^Department of Basic Medical Sciences, ^2^Department of Medicinal Chemistry and Molecular Pharmacology, ^3^Purdue Center for Cancer Research, Purdue University, and ^4^Gene Editing Core, Bindley Biosciences Center, Purdue University, West Lafayette, IN 47907. ^5^Shanghai Institute of Biochemistry and Cell Biology, Chinese Academy of Sciences, Suzhou, Jiangsu 215121, China. ^6^Department of Hepatology, Hôpital de la Croix-Rousse, Hospices Civils de Lyon, Université Lyon 1, Lyon, France

Department of Basic Medical Sciences,

Purdue University

201 S. University Street

West Lafayette, IN  47907-2064

**Supplementary Figures**

**Figure S1**
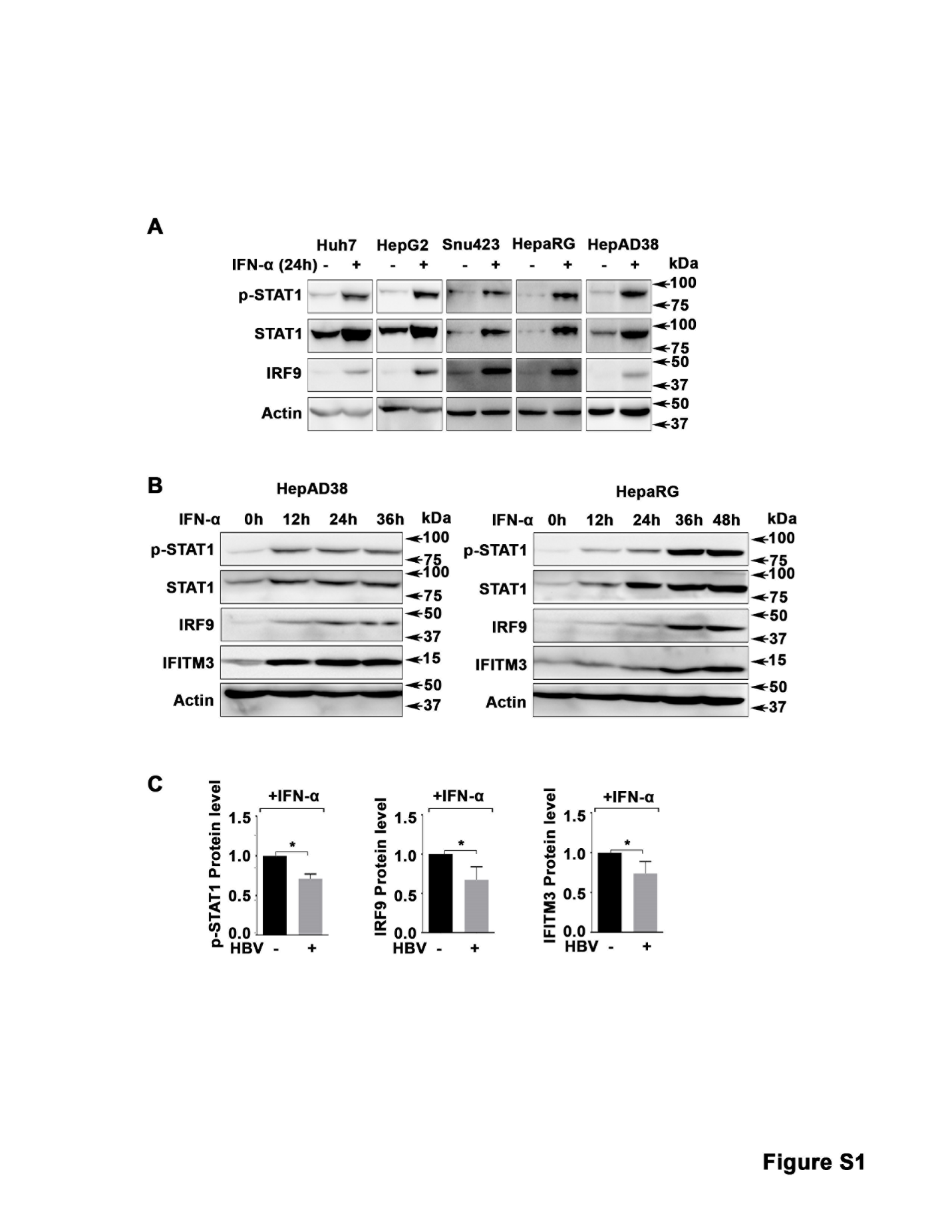

**Figure S1. IFN-α response in various liver cancer cell lines.** Immunoblots of IFN-α signaling related proteins using lysates from **(A)** Huh7, HepG2, Snu423, HepaRG and HepAD38 cells 24 h after IFN-α (500 ng/ml) treatment; **(B)** time course, 12-36 h of IFN-α (500 ng/ml) treatment in HepAD38 and HepaRG cells. **(C)** Quantification of indicated proteins from immunoblots in Figure 1A, using imageJ software**.**

**Figure S2**

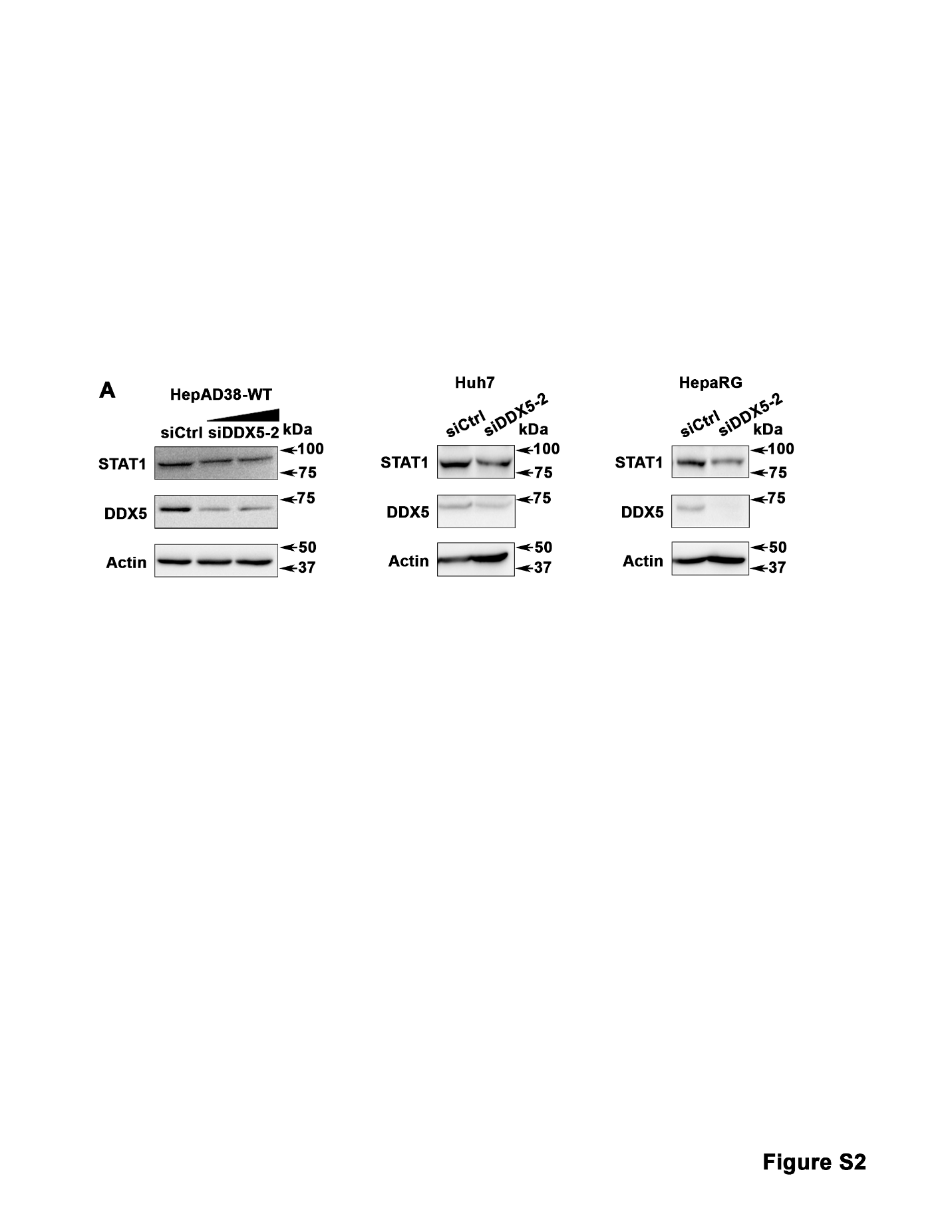

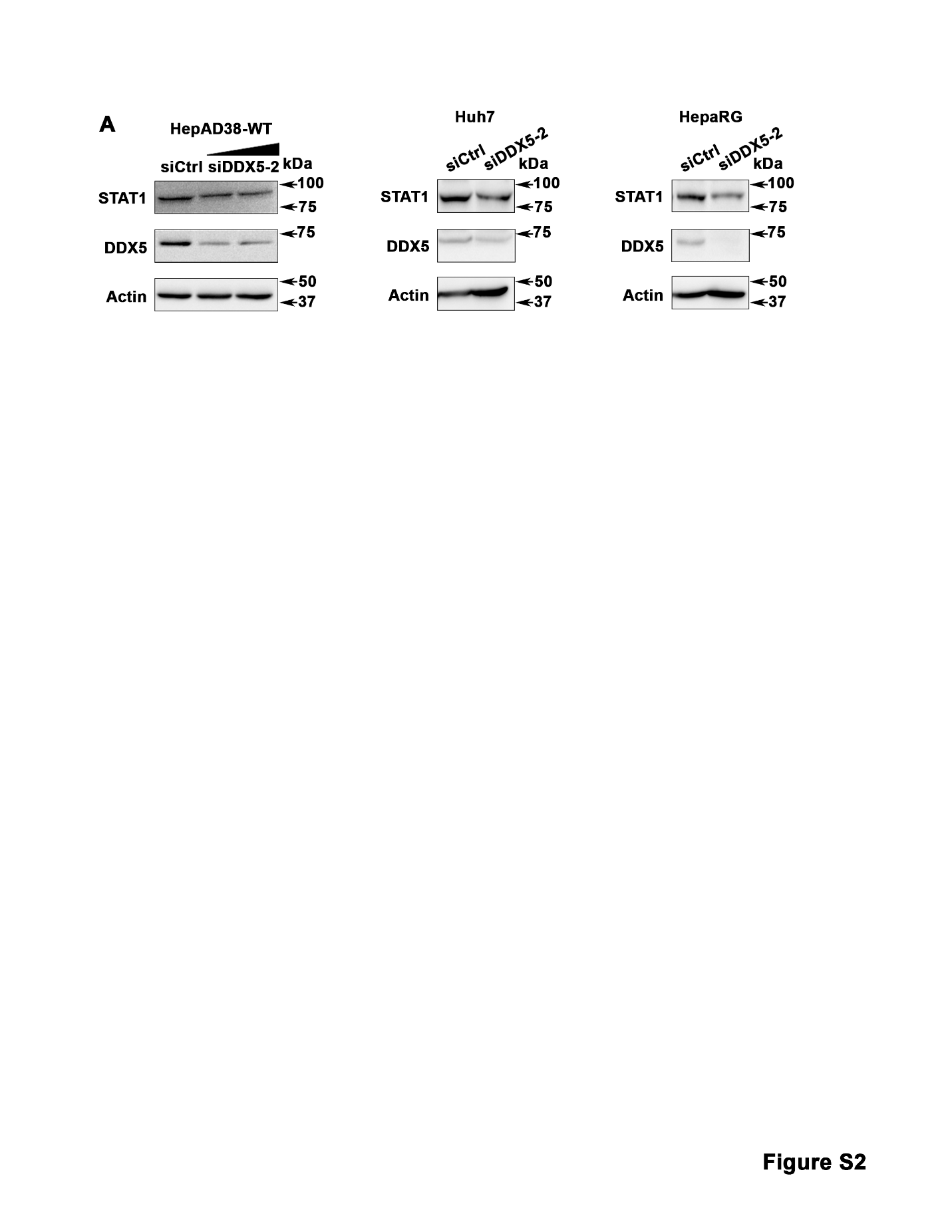
**Figure S2. DDX5 knockdown reduces STAT1 protein level.** Immunoblots of indicated proteins, in HepAD38, Huh7 and HepaRG cells using another sequence for siRNA for DDX5, indicated as siDDX5-2.

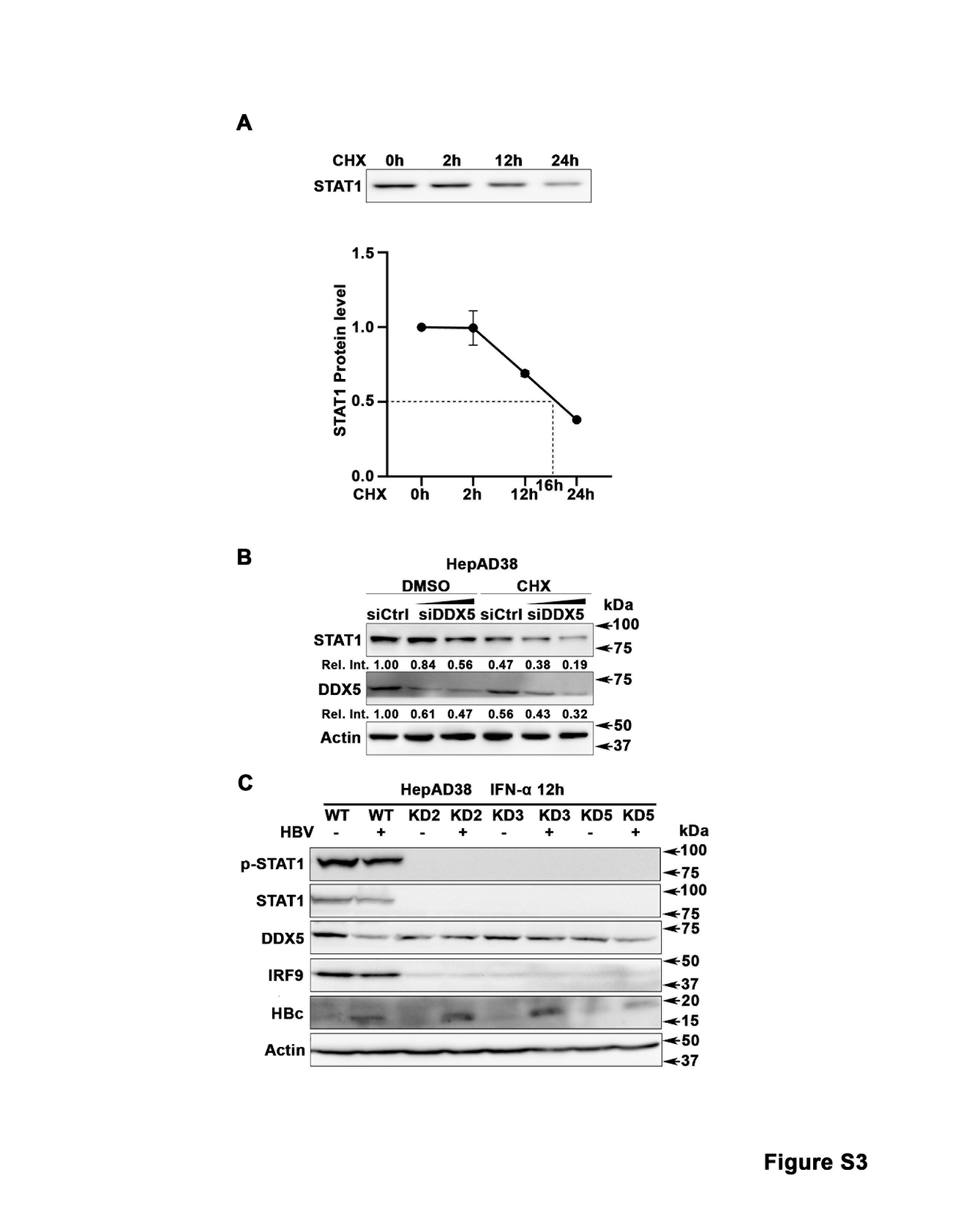
**Figure S3**

**Figure S3. Quantification of t1/2 of STAT1 protein. (A)** Immunoblot of STAT1 following treatment with 20 µg/ml cyclohexamide (CHX) for indicated time course in HepAD38 cells. Graph shows quantification of STAT1 t1/2. **(B)** Immunoblot of indicated proteins using lysates from HepAD38 treated or not with CHX **(**20 µg/ml) for 12h and transfected with siCtrl or increasing amount (20-40 nM) siDDX5 RNA. **(C)** Immunoblot of lysates from WT and KD2, KD3 and KD5 HepAD38 cells, treated with IFN-α for 12 h, +/- HBV replication for 3 days, for indicated proteins.

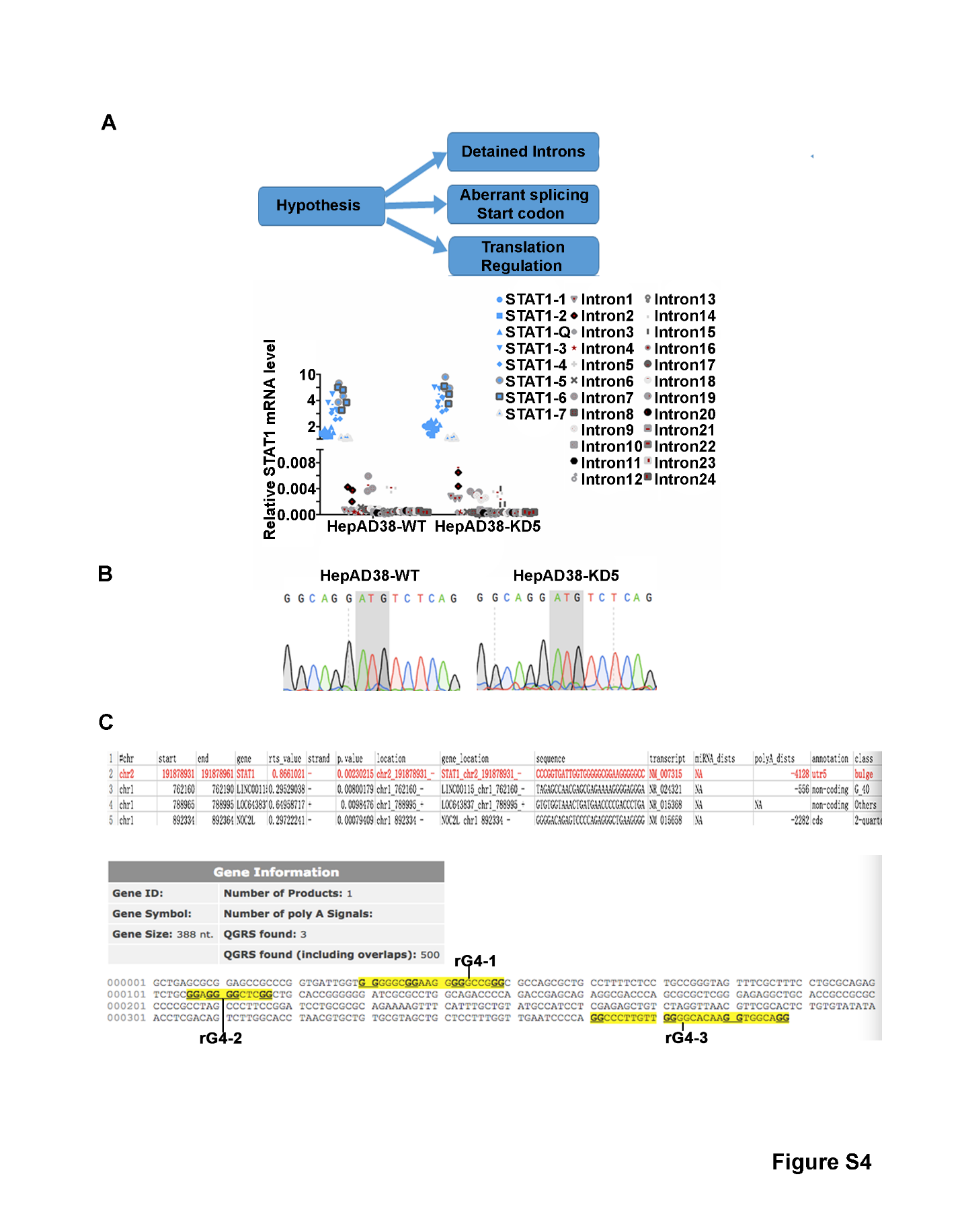
**Figure S4**

**Figure S4. Mechanism of DDX5 effect on STAT1 regulation. (A)** Diagram of post-transcriptional effects of DDX5 on STAT1 expression. QRT PCR quantifying expression of indicated exons (blue) and indicated introns (gray) in HepAD38-WT and HepAD38-KD5 (DDX5-knockdown) cell lines. N=3. Primer sequences are in Supplementary Table S3**. (B)** DNA sequencing of 5’UTR of STAT1 mRNA isolated from HepAD38-WT or HepAD38-KD5 cells. **(C).** Table listing the 5’UTR of STAT1 mRNA as containing a high probability G-quadruplex Structure (1)**.** Nucleotide sequence corresponding to the 5’UTR of human STAT1 mRNA.The highlighted sequences indicate the putative rG4 sequences.

**Figure S5**

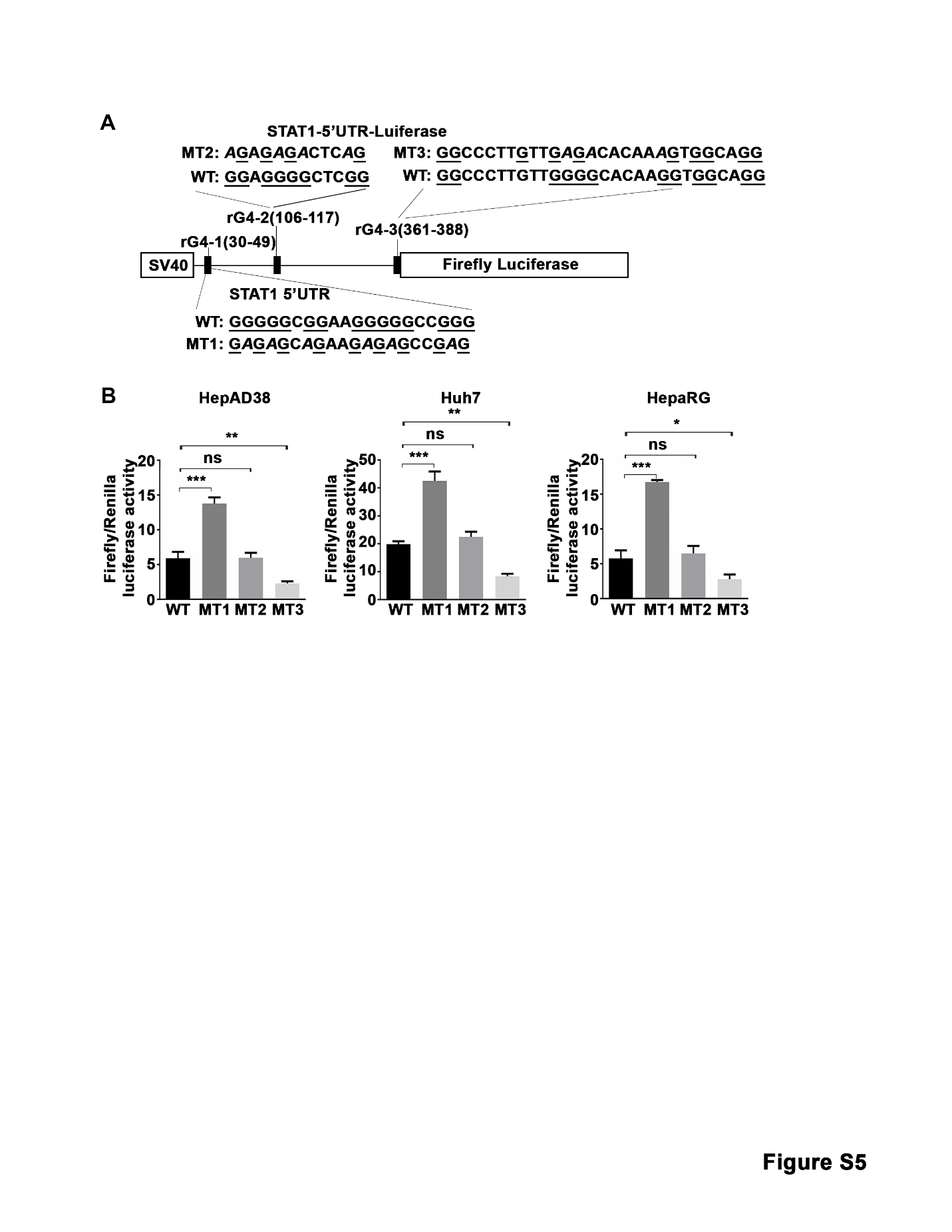

**Figure S5. Putative rG4-2 and rG4-3 sequences of STAT1 5’UTR do not affect STAT1 translation. (A)** Human STAT1 5’UTR upstream of Firefly (F.) Luciferase reporter, driven from the SV40 promoter. Putative rG4 sequences in 5’ UTR indicated as rG4-1, rG4-2 and rG4-3. WT rG4-1 sequence is shown. Italics indicate site-directed changes in mutant MT1 of rG4-1, MT2 of rG4-2 and MT3, of rG4-3. **(B)** Ratio of Firefly/Renilla Luciferase activity at 24 h after co-transfection of WT or MT1, MT2 and MT3 STAT1-5’UTR-Luciferase and Renilla-Luciferase expression plasmids, in indicated cell lines. Statistical analysis from three independent biological replicates. *: p<0.05, **: p<0.01, ***: p<0.001, ns: not significant; Error bars indicate Mean ±SEM.

**Figure S6**
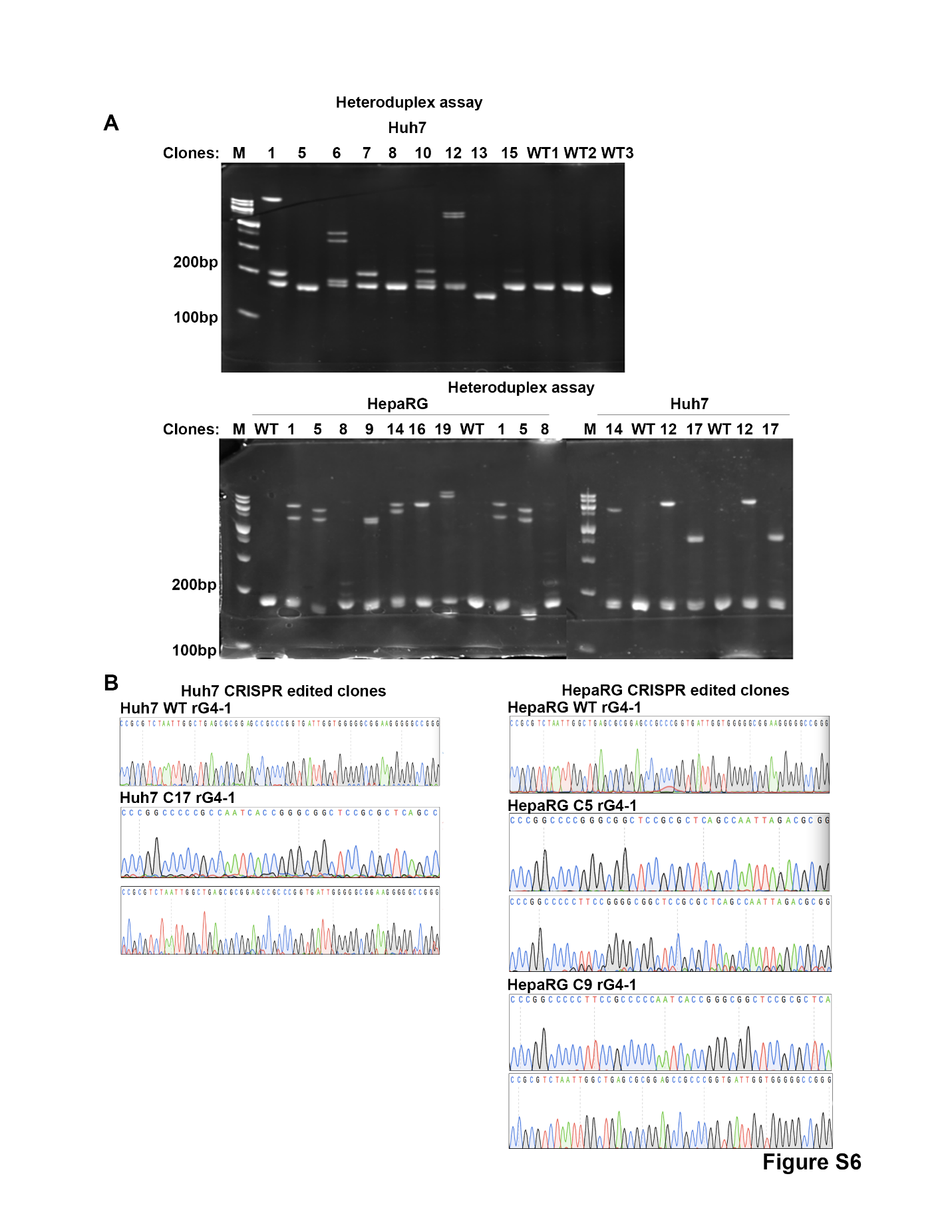

**Figure S6**. **Analyses of CRISPR/Cas9 edited cell lines:** **(A)** heteroduplex assay of various edited clones by agarose gel electrophoresis. **(B)** DNA sequence analyses of indicated edited clones.

**Figure S7**

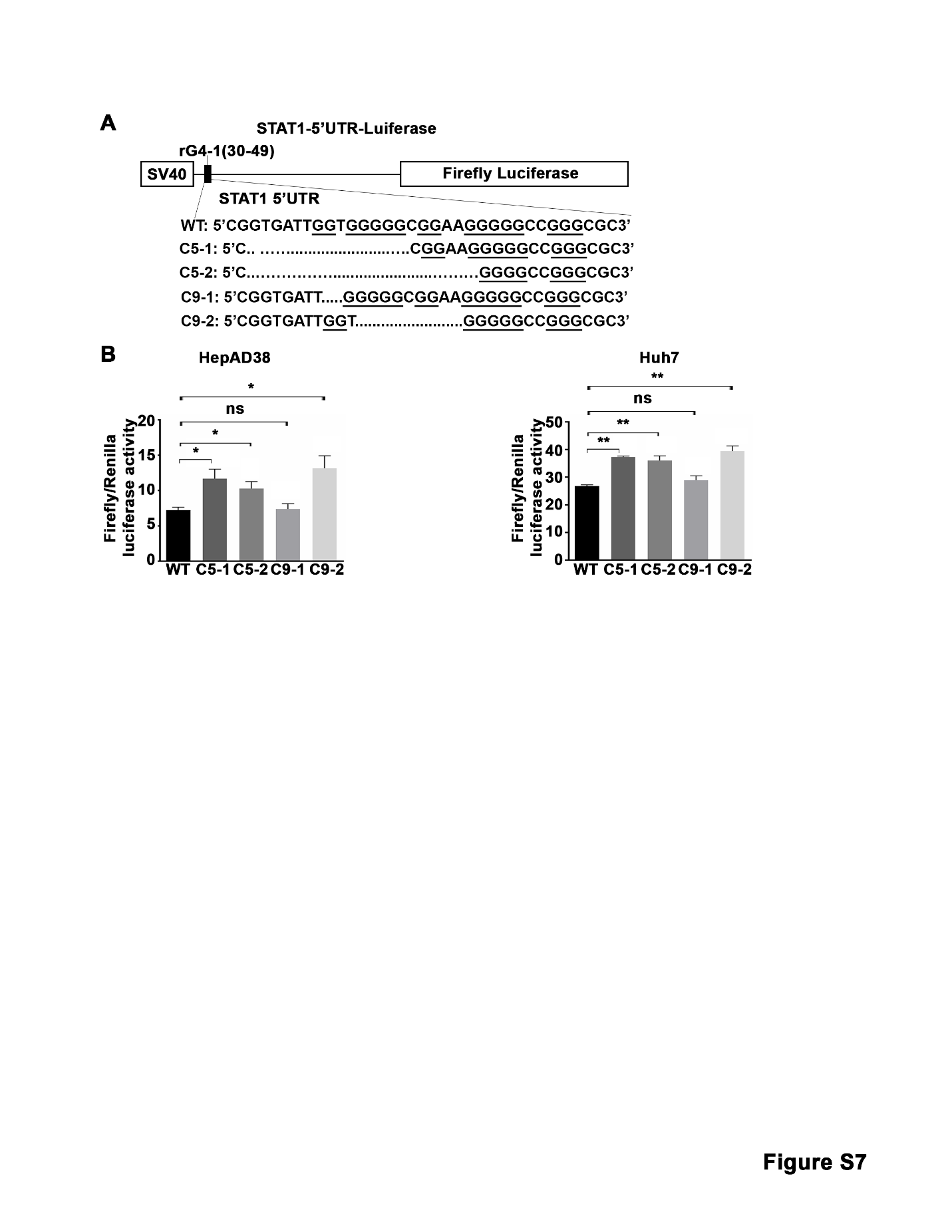

**Figure S7. Activity of edited rG4-1 sequence studied by Luciferase reporter assays. (A)** Sequence of edited rG4-1 sequences cloned into the indicated Luciferase expression vector. **(B) (B)** Ratio of Firefly/Renilla Luciferase activity at 24 h after co-transfection of WT and edited rG4-1 sequences in STAT1-5’UTR-Luciferase and Renilla-Luciferase expression plasmids, in indicated cell lines. Statistical analysis from three independent biological replicates. *: p<0.05, **: p<0.01, ***: p<0.001, ns: not significant; Error bars indicate Mean ±SEM.

**Figure S8.**
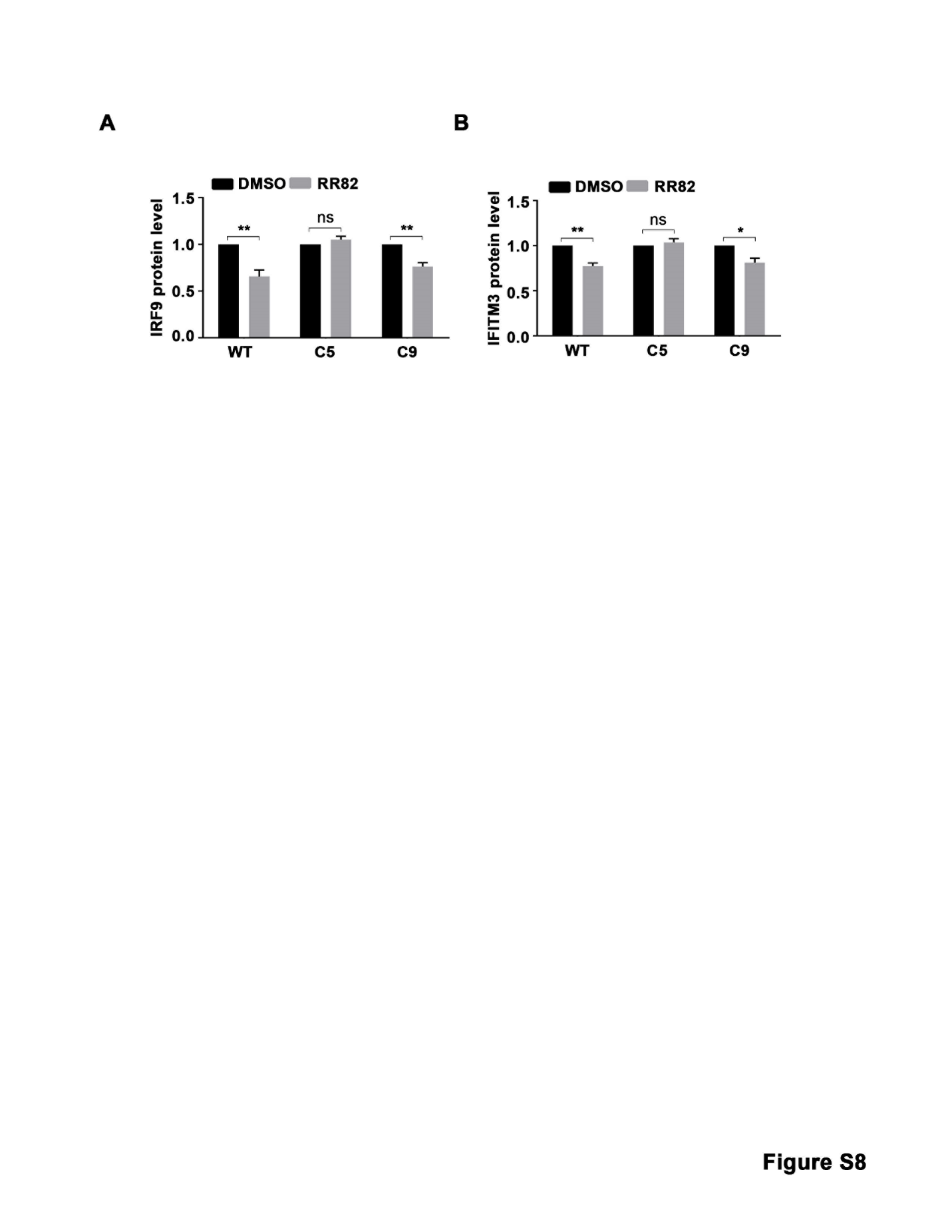

**Figure S8.** Quantification of immunoblots from Figure 5C using imageJ software. N=3

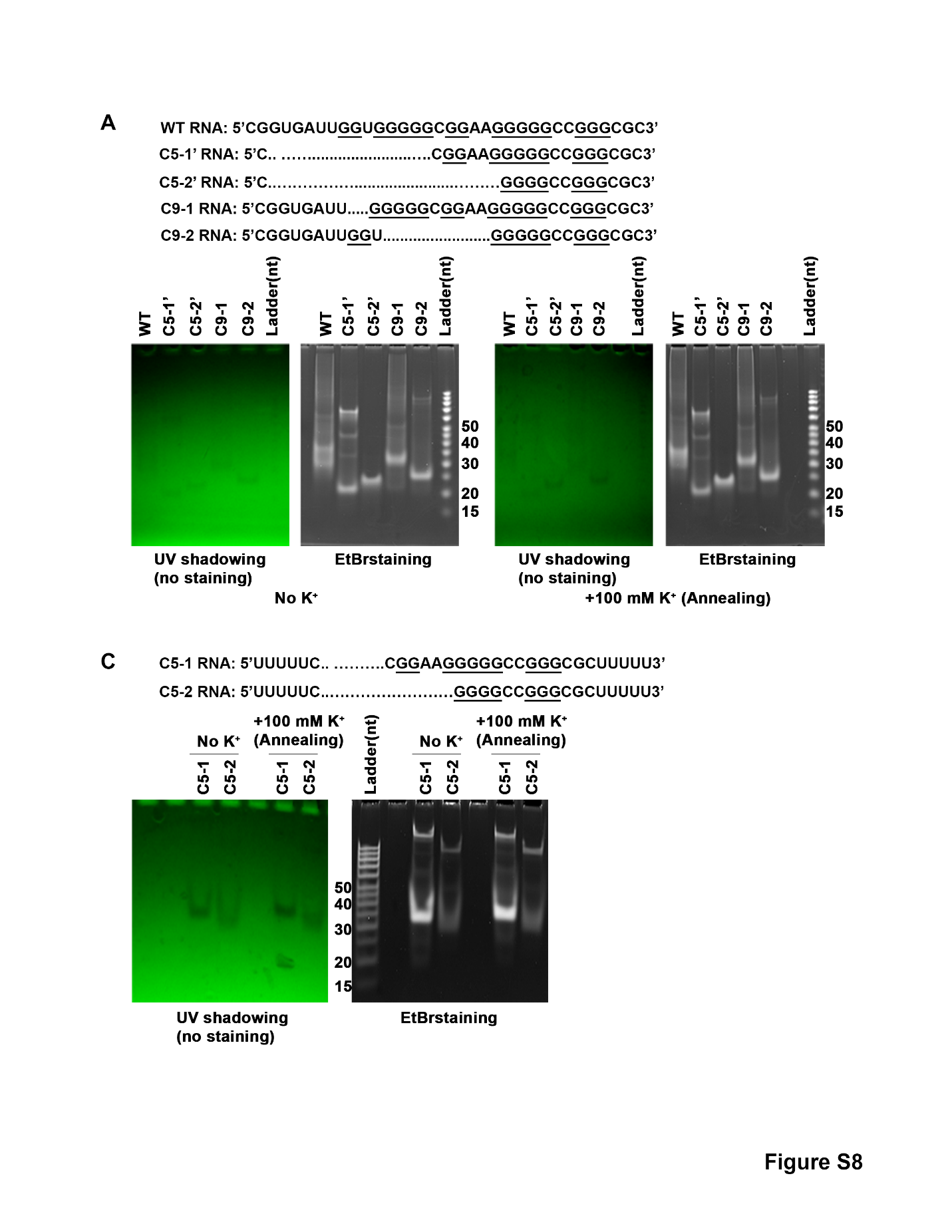
**Figure S9**

**Figure S9. Analyses of synthesized RNA oligonucleotides** shown in **(A)** by native agarose gel electrophoresis **(B and C).**

**Figure S10**

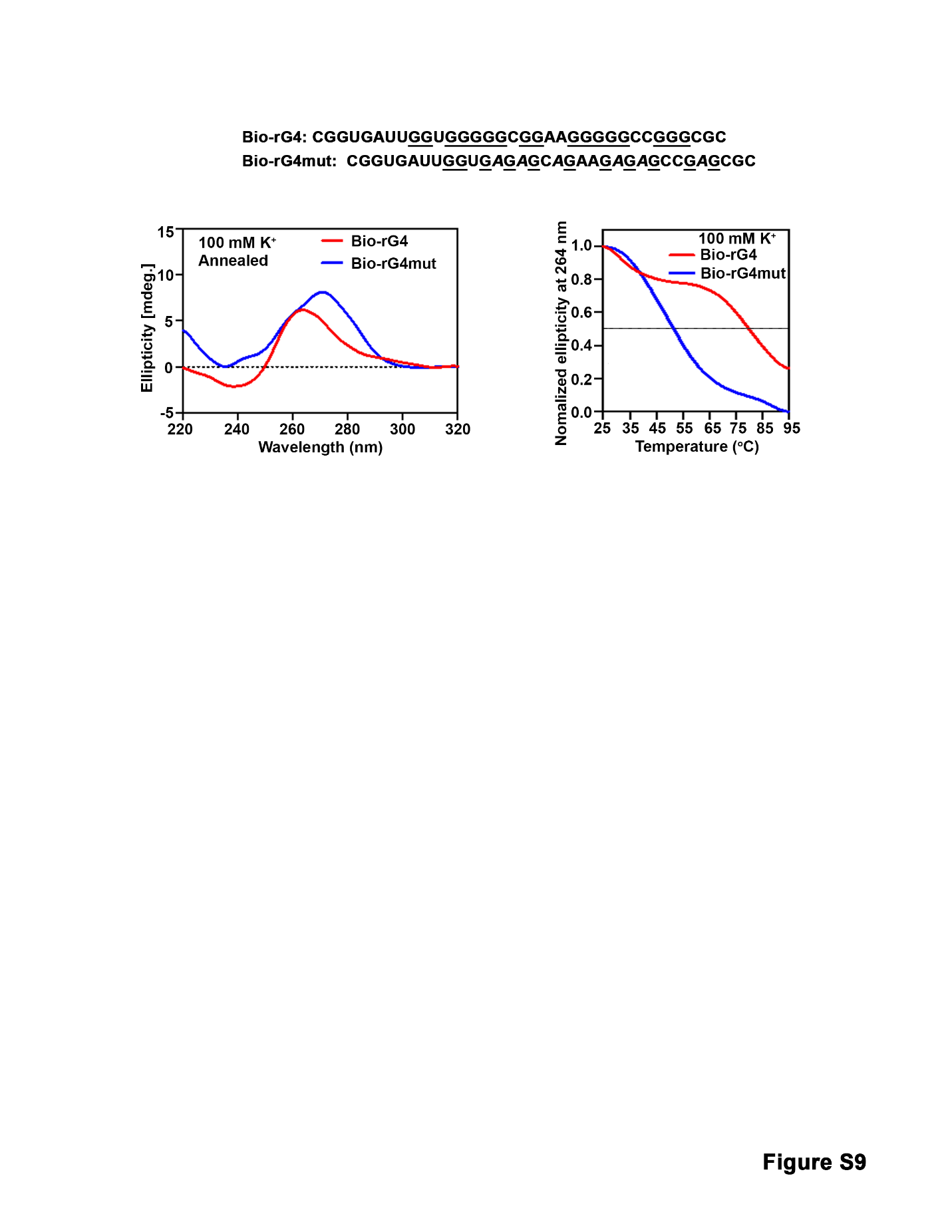

**Figure S10. CD spectroscopy and melting curves of** **synthetic biotinylated RNA oligonucleotides** **used in pulldown assays**. The sequence of bio-rG4 and bio-rG4mut of STAT1 is shown; Analyses included CD spectroscopy measurements (left panel) and melting curves (right panel). RNA oligonucleotides annealed by heating to 95^o^C and slowly cooled down to room temperature, in the presence of 100 mM K^+^.

**Figure S11**

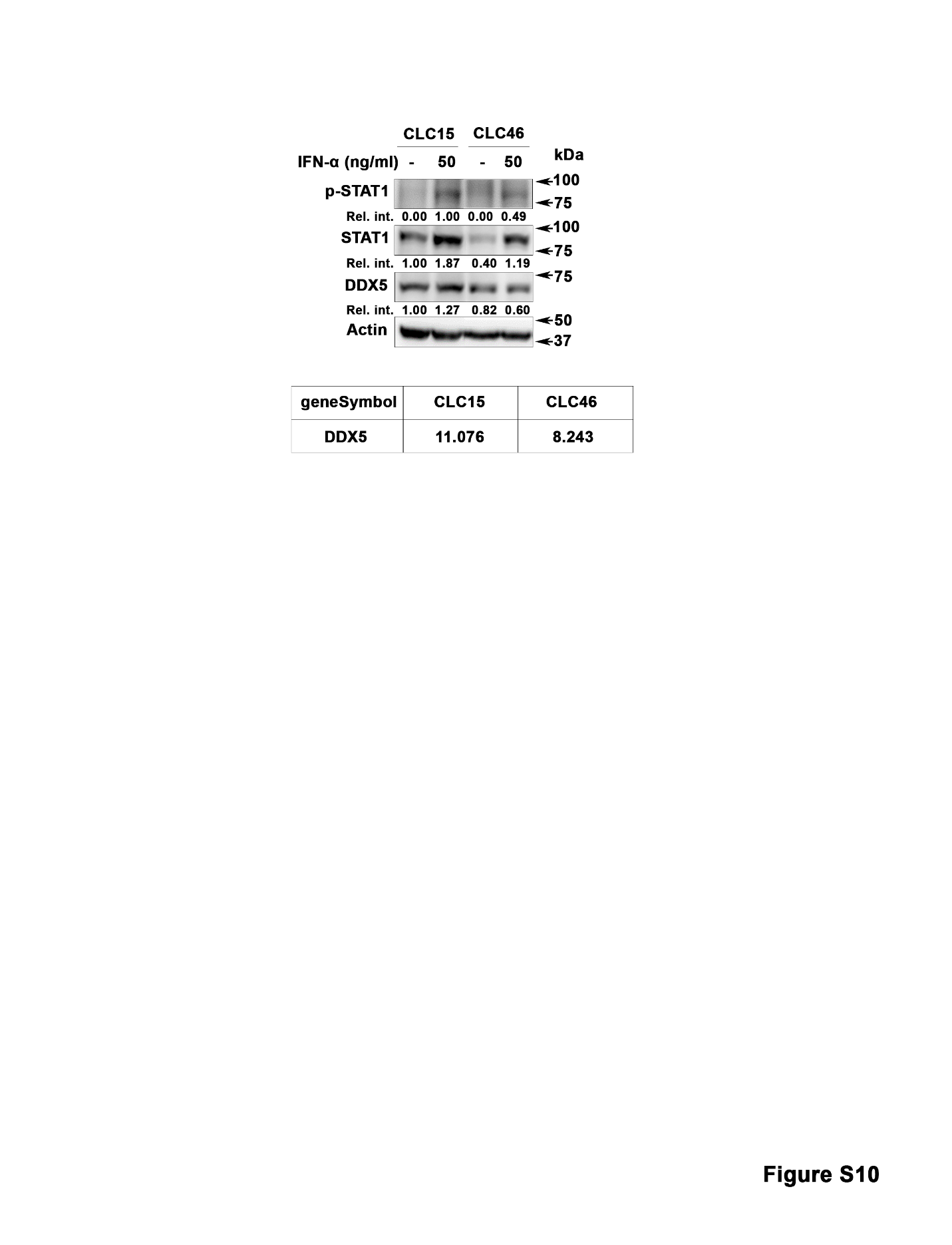

**Figure S11.** Immunoblots using lysates of HBV-HCC derived cell lines (28) treated with IFN-α (50 ng/ml) for 12 h. Quantification by image J software is relative to actin. A representative assay is shown, n=2. The log_2_ RPKM values of DDX5 in CCL15 and CCL46 cell lines from the RNAseq studies (28) is shown.

**Supporting Table S1: List of Plasmids and siRNAs**

| **Plasmids, siRNAs** | **Source** |
| --- | --- |
| Wild Type (WT) STAT1-5’UTR-Luciferase | Addgene (#115353)  (pFL-SV40-STAT1 5'-UTR was a gift from Ming-Chih Lai) |
| Mutant (MT)-STAT1-5’UTR-Luciferase | Constructed by site directed mutagenesis using the full-length  WT 5’UTR as a template, and QuikChange II XL Site-Directed Mutagenesis kit (Agilent). |
| Renilla luciferase vector | Addgene (#27163) |
| siCtrl | ThermoFisher Scientific (#4390843) |
| siDDX5-1 | ThermoFisher Scientific (#4392420, assay id s4007) |
| siDDX5-2 | ThermoFisher Scientific (#4392420, assay id s4008) |

**Supporting Table S2: Antibodies**

| **Antibody** | **Dilution** | **Application** | **Source** |
| --- | --- | --- | --- |
| Mouse α-Human p68 | 1:1000 in 2% BSA in TBST | Western Blot | Millipore Sigma (#05-850) |
| Rabbit α-Human STAT1 | 1:1000 in 2% BSA in TBST | Western Blot | Cell Signaling Technologies (#14994S) |
| Rabbit α-HBV Core | 1:5000 in 2% BSA in TBST | Western Blot | Dr. Adam Zlotnick lab |
| Mouse α-Human Actin | 1:1000 in 2% BSA in TBST | Western Blot | Sigma (#A5441) |
| Rabbit α-Human p-STAT1 | 1:1000 in 2% BSA in TBST | Western Blot | Millipore Sigma (#07-307) |
| Rabbit α-Human IRF9 | 1:1000 in 2% BSA in TBST | Western Blot | Cell Signaling Technologies (#76684S) |
| Rabbit α-Human IFITM3 | 1:1000 in 2% BSA in TBST | Western Blot | Cell Signaling Technologies (#59212S) |
| Rabbit α-Human NRAS | 1:1000 in 2% BSA in TBST | Western Blot | Proteintech (#10724-1-AP) |
| Horse α-Mouse secondary | 1:2000 in 2% BSA in TBST | Western Blot | Vector Laboratories (#PI-2000) |
| Goat α-Rabbit secondary | 1:2000 in 2% BSA in TBST | Western Blot | Vector Laboratories (#PI-1000) |
| Rabbit α-Human DDX5 | 5 μg | RNA Immunoprecipitation | Cell Signaling Technologies (#9877S) |
| Rabbit IgG | 5 μg | RNA Immunoprecipitation | Millipore Sigma (#17-700) |
| Rabbit α-Human DDX5 | 1:1000 in 2% BSA in TBST | Immunohistochemistry | Abacm (#ab205718) |

**Supporting Table S3: Primer and RNA oligonucleotide sequences**

| **Primer** | **5’ – Sequence – 3’** |
| --- | --- |
| STAT1-Q-F | ATCAGGCTCAGTCGGGGAATA |
| STAT1-Q-R | TGGTCTCGTGTTCTCTGTTCT |
| DDX5-F | AGCAAGTGAGCGACCTTATC |
| DDX5-R | CATCCTTCATGCCTCCTCTAC |
| GAPDH-F | CCCTTCATTGACCTCAACTACA |
| GAPDH-R | ATGACAAGCTTCCCGTTCTC |
| Actin-F | GGCATGGGTCAGAAGGATT |
| Actin-R | GGGGTGTTGAAGGTCTCAAA |
| UBc-F | CCTGGAGGAGAAGAGGAAAGAGA |
| UBc-R | TTGAGGACCTCTGTGTATTTGTCA |
| HBV pgRNA-F | CTCCTCCAGCTTATAGACC |
| HBV pgRNA-R | GTGAGTGGGCCTACAAA |
| STAT1-1F | GCGCGCAGAAAAGTTTCATTTGC |
| STAT1-1R | CTGAGACATCCTGCCACCTTG |
| STAT1-2F | GTACGAACTTCAGCAGCTTGAC |
| STAT1-2R | GAAAATTATCCTGAAGATTACGC |
| STAT1-3F | AACGGAGGCGAACCTGAC |
| STAT1-3R | AACGGAGGCGAACCTGAC |
| STAT1-4F | AGAGCCAATGGAACTTGATGG |
| STAT1-4R | ACTATCCGAGACACCTCGTC |
| STAT1-5F | TCCTGCTACTCTGTTCCTTCAC |
| STAT1-5R | CCCTCATTCTCGTCCTGATAC |
| STAT1-6F | CTGCTTTCATCTTGGTCACATAC |
| STAT1-6R | AAAGTAGCCCATTTAAGAAACATG |
| STAT1-7F | ATGGCGAGAACCTAAGTTTCAG |
| STAT1-7R | GCAGTAAAATGAAACCATGCCG |
| Intron-1F | GCGCGCAGAGTGAGTGGCCG |
| Intron-1R | TGGAATACTCAGGACGCGTTC |
| Intron-2F | CAAGGTGGCAGGTCAGTAAATG |
| Intron-2R | GAACTCTTCAGTACCGGCCAG |
| Intron-3F | CAAGACTGGTAAGGAAAATTCAC |
| Intron-3R | GTTGAGACACTTTTTGGCTCAAC |
| Intron-4F | CGTAATCTTCAGGTATGACCTGG |
| Intron-4R | ATGCTATATTTACTGATGCTGTAG |
| Intron-5F | AATCAGGTACTTTTTTCTCATTG |
| Intron-5R | CTCTAAATCTGATTCTCCCACTTC |
| Intron-6F | CAAGGTTATGGTGAGTATTGAG |
| Intron-6R | ACTTATAGCTTGAGACTTCTGC |
| Intron-7F | AGAACAGAGGTAAGGGTTCAC |
| Intron-7R | TGGCCTGGGTTATCAAGGAAG |
| Intron-8F | GAGAAAGGTAGTTATTTACTTTCC |
| Intron-8R | TTTCATTCTCAACTGGGGCTC |
| Intron-9F | GAACTGGTAAGATTCTCCAAAGC |
| Intron-9R | CTGTGCTTTCCAAGGGAGTG |
| Intron-10F | ATTCAGAGGTAACTCAAGGGAC |
| Intron-10R | CCTGGGTGATAGGTGAGACTC |
| Intron-11F | GTTGAGGTAACAAGGGAAAGATG |
| Intron-11R | CTCTCAGATATTCTCAGTAAGAG |
| Intron-12F | ATTTGATAAGTAAGATATCTTTAAC |
| Intron-12R | GGCAAACTTCCACCCAGTATAG |
| Intron-13F | ACAGTAAAAGGGTACGTGACG |
| Intron-13R | CTATACAATATAGGAAAGAAATGC |
| Intron-14F | GGCACCTGGTAGGGACATCA |
| Intron-14R | AGTACTGGCGACAGGAAGAC |
| Intron-15F | CCAGAACGAATGAGGTGAGAG |
| Intron-15R | GTGGTGTTTTTACTGTTGTCACAG |
| Intron-16F | GGTAATTGACCTCGAGGTAAGAC |
| Intron-16R | CAATTGTCTAGCTTTCTGGACCA |
| Intron-17F | AACCCAGGGTATGGAAAACAC |
| Intron-17R | ATATCCATGCTTCATATTCCCAG |
| Intron-18F | GAGAGAAGCTTCTTGGTATATGC |
| Intron-18R | GTGACATATGATTCTCACTTAGC |
| Intron-19F | GGTTTTGTAAGGTGAGGACTG |
| Intron-19R | AATGTTATCACTCAGGCAGGTG |
| Intron-20F | GGAATGATGGGTAAGGGCCA |
| Intron-20R | GTTCTACTCTTCTGAAGCCCTG |
| Intron-21F | AGGCGGTGAGTGGGAGTTTG |
| Intron-21R | AGAGAGTGAACCACAACCAC |
| Intron-22F | TCCAGGCCAAAGGAAGGTAAG |
| Intron-22R | GCTGTATCAGGCCAAATAATGC |
| Intron-23F | GTGTCTGAAGTGTAAGTGAACAC |
| Intron-23R | AAATGCTGATAGGCAGTAACAC |
| Intron-24F | CGACAGTATGGTGAGTACCAC |
| Intron-24R | AGGTGGTTTAAATCCAGCAGC |
| Fluc-F | CTCACTGAGACTACATCAGC |
| Fluc-R | TCCAGATCCACAACCTTCGC |
| Rluc-F | GGAATTATAATGCTTATCTACGTGC |
| Rluc-R | CTTGCGAAAAATGAAGACCTTTTAC |
| STAT1-F 5'UTR EcoRI | CGGAATTCGCTGAGCGCGGAGCCGCCCGG |
| Stat1-F 5'UTR NcoI | CATGCCATGGCCTGCCACCTTGTGCCCCAAC |
| MT1-F | CCGGTGATTGGTGAGAGCAGAAGAGAGCCGAGCGCCAGCGCTGC |
| MT1-R | GCAGCGCTGGCGCTCGGCTCTCTTCTGCTCTCACCAATCACCGG |
| MT2-F | GCGCAGAGTCTGCAGAGAGACTCAGCTGCACCGGGGG |
| MT2-R | CCCCCGGTGCAGCTGAGTCTCTCTGCAGACTCTGCGC |
| MT3-F | GAATCCCCAGGCCCTTGTTGAGACACAAAGTGGCAGGCCATGGAAGACG |
| MT3-R | CGTCTTCCATGGCCTGCCACTTTGTGTCTCAACAAGGGCCTGGGGATTC |
| MT-C5-1-F | GCTGAGCGCGGAGCCGCCCCGGAAGGGGGCCGGGCGCCA |
| MT-C5-1-R | TGGCGCCCGGCCCCCTTCCGGGGCGGCTCCGCGCTCAGC |
| MT-C5-2-F | GCTGAGCGCGGAGCCGCCCGGGGCCGGGCGCCAGCGCTG |
| MT-C5-2-R | CAGCGCTGGCGCCCGGCCCCGGGCGGCTCCGCGCTCAGC |
| MT-C9-1-F | CGCGGAGCCGCCCGGTGATTGGGGGCGGAAGGGGGCCGGG |
| MT-C9-1-R | CCCGGCCCCCTTCCGCCCCCAATCACCGGGCGGCTCCGCG |
| MT-C9-2-F | GGAGCCGCCCGGTGATTGGTGGGGGCCGGGCGCCAGCGC |
| MT-C9-2-R | GCGCTGGCGCCCGGCCCCCACCAATCACCGGGCGGCTCC |
| STAT1-seq-F | TGTATGCCATCCTCGAGAGC |
| STAT1-seq-R | CAGTCTTGCTTTTCTAACCACTG |
| SV40-F | TATTTATGCAGAGGCCGAGG |
| M13-R | CAGGAAACAGCTATGAC |
| CRISPR-seq-F | AAGCCGGCGGAAATACCC |
| CRISPR-seq-R | TCTGCTCGGTCTGGGGTC |
| CRISPR-heter-F | GGAAGCCGGCGGAAATAC |
| CRISPR-heter-R | GCGCAGGAAAGCGAAACTA |
| CRISPR-gRNA1 | rArGrCrCrGrCrCrCrGrGrUrGrArUrUrGrGrUrGrG |
| CRISPR-gRNA2 | rCrGrGrUrGrArUrUrGrGrUrGrGrGrGrGrCrGrGrA |
| WT (rG4-1) RNA oligos | rCrGrGrUrGrArUrUrGrGrUrGrGrGrGrGrCrGrGrArArGrGrGrGrGrCrCrGrGrGrCrGrC |
| C5-1 RNA oligos | rUrUrUrUrUrCrCrGrGrArArGrGrGrGrGrCrCrGrGrGrCrGrCrUrUrUrUrU |
| C5-2 RNA oligos | rUrUrUrUrUrCrGrGrGrGrCrCrGrGrGrCrGrCrUrUrUrUrU |
| C9-1 RNA oligos | rCrGrGrUrGrArUrUrGrGrGrGrGrCrGrGrArArGrGrGrGrGrCrCrGrGrGrCrGrC |
| C9-2 RNA oligos | rCrGrGrUrGrArUrUrGrGrUrGrGrGrGrGrCrCrGrGrGrCrGrC |
| C5-1’ RNA oligos | rCrCrGrGrArArGrGrGrGrGrCrCrGrGrGrCrGrC |
| C5-2’ RNA oligos | rCrGrGrGrGrCrCrGrGrGrCrGrC |
| Bio-rG4 RNA oligos | rCrGrGrUrGrArUrUrGrGrUrGrGrGrGrGrCrGrGrArArGrGrGrGrGrCrCrGrGrGrCrGrC |
| Bio-rG4mut RNA oligos | rCrGrGrUrGrArUrUrGrGrUrGrArGrArGrCrArGrArArGrArGrArGrCrCrGrArGrCrGrC |

**Supporting Table S4: Reagents, Chemical inhibitors, and Kits**

| **Reagents, Chemical inhibitors, Kits** | **Source** |
| --- | --- |
| Cycloheximide (CHX) | Cell Signaling Technologies (#2112) |
| IFN-α | Millipore Sigma (#SRP4596) |
| RR82 | ChemSpider (324571151) |
| PhenDC3 | Polysciences (#26000-1) |
| TMPyP2 | Dr. Danzhou Yang |
| TMPyP4 | Dr. Danzhou Yang |
| PCR Mycoplasma Detection Kit | Abm (#G238) |
| Dual-Luciferase® Reporter Assay System | Promega (#1980) |
| Cell Lysis Buffer (10X) | Cell Signaling Technology (#9803) |
| LightCycler® 480 SYBR Green I Master | Roche (#04887352001) |
| iScript™ cDNA Synthesis Kit | Biorad (#1708891) |
| Nitrocellulose Membrane, Roll, 0.2 µm | Biorad (#1620112) |
| LightCycler® 480 Sealing Foil | Roche (#04729757001) |
| LightCycler® 8-Tube Strips (white) | Roche (#06612601001) |
| DMSO | Sigma (#D8418-50ML) |
| Tween™ 20 | ThermoFisher Scientific (BP337-500) |
| Tetracycline hydrochloride | Sigma (#T7660-5G) |
| Bovine Serum Albumin | Sigma (#A9647-100G) |
| Pierce™ ECL Western Blotting Substrate | ThermoFisher Scientific (#32106) |
| Geneticin™ Selective Antibiotic (G418 Sulfate) | ThermoFisher Scientific (#10131027) |
| Pierce™ BCA Protein Assay Kit | ThermoFisher Scientific (#23227) |
| Lipofectamine™ 3000 Transfection Reagent | ThermoFisher Scientific (#L3000015) |
| Lipofectamine™ RNAiMAX Transfection Reagent | ThermoFisher Scientific (#13778150) |
| Restore™ PLUS Western Blot Stripping Buffer | ThermoFisher Scientific (#46430) |
| RNeasy Mini Kit | Qiagen (#74104) |
| Magna RIP™ RNA-Binding Protein Immunoprecipitation Kit | Millipore Sigma (#17-700) |
| CRISPRevolution sgRNA EZ Kit | SYNTHEGO |
| QuikChange II XL Site-Directed Mutagenesis Kit | Agilent (#200521) |
| Streptavidin MagneSphere Paramagnetic Particles | Promega (#Z5481) |
| Glycerol | Promega (#H5433) |
| KCl (2M), RNase-free | Fisher Scientific (AM9640G) |
| LiCl Precipitation Solution (7.5 M) | Fisher Scientific (AM9480) |
| RIPA Lysis and Extraction Buffer | Fisher Scientific (89900) |
